# Neuroanatomical Norms in the UK Biobank: The Impact of Allometric Scaling, Sex and Age

**DOI:** 10.1101/2020.12.14.422684

**Authors:** Camille Michèle Williams, Hugo Peyre, Roberto Toro, Franck Ramus

## Abstract

Few neuroimaging studies are sufficiently large to adequately describe population-wide variations. This study’s primary aim was to generate neuroanatomical norms and individual markers that consider age, sex, and brain size, from 629 cerebral measures in the UK Biobank (N = 40 028). The secondary aim was to examine the effects and interactions of sex, age, and brain allometry – the non-linear scaling relationship between a region and brain size (e.g., Total Brain Volume) across cerebral measures.

Allometry was a common property of brain volumes, thicknesses, and surface areas (83%) and was largely stable across age and sex. Sex differences occurred in 67% of cerebral measures (median |β|= 0.13): 37% of regions were larger in males and 30% in females. Brain measures (49%) generally decreased with age, although aging effects varied across regions and sexes. While models with an allometric or linear covariate adjustment for brain size yielded similar significant effects, omitting brain allometry influenced reported sex differences in variance.

This large scale-study advances our understanding of age, sex, and brain allometry’s impact on brain structure and provides data for future UK Biobank studies to identify the cerebral regions that covary with specific phenotypes, independently of sex, age, and brain size.

**Graphical Abstract:** 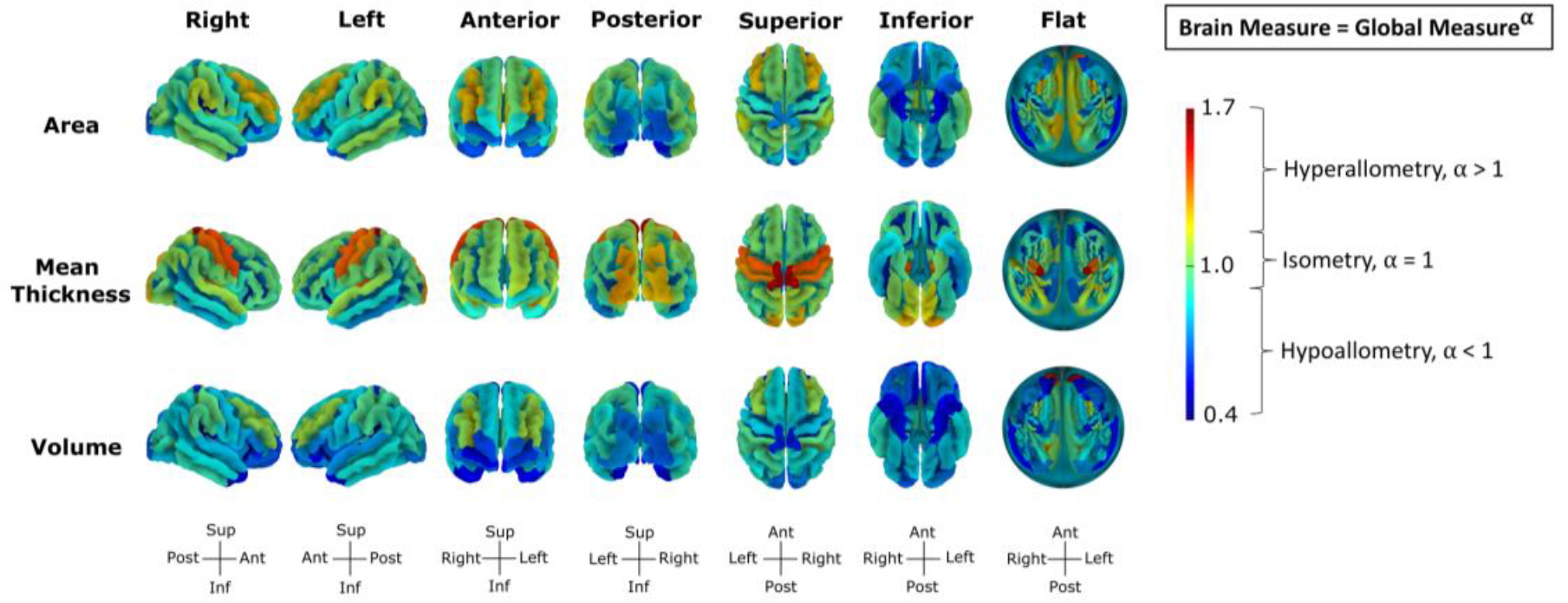

**Highlights:**

- We created neuroanatomical norms and individual markers for the UK Biobank (N=40 028)
- Allometry was common across 83% of brain volumes, thicknesses, and surface areas
- 67% of regions differed between sexes: 37% were larger in males and 30% in females
- Omitting brain allometry influenced reported sex differences in variance
- 49% of regions declined with age, with variations across regions and sexes

## 1. Introduction

Although all humans share a common brain structure and organization, they also vary in terms of the size and shape of their brain and its subcomponents. These neuroanatomical variations are thought to partly underlie differences in cognitive and behavioral traits and in the risk of developing psychiatric and neurological disorders (for review Dallaire-Théroux et al., 2017; Deary, 2010; Jumah et al., 2016; Oakes et al., 2017; Schmidt et al., 2018). Yet, most of these studies rely on relatively small samples and suffer from high sampling variability. When the sample is too small to accurately represent the control or target population, spurious neuroanatomical markers may be reported. Moreover, if true effects are observed in small samples, their size would be exaggerated as only large effects would pass a conventional statistical significance threshold (e.g., p < 0.05) with few degrees of freedom (for review Szucs & Ioannidis, 2017, 2020). Thus, despite hundreds of studies, few neuroanatomical measures can be declared as robust markers of cognitive traits or psychiatric and neurological disorders (Gong et al., 2019; Marek et al., 2020; Matsuo et al., 2019; Peyre et al., 2020; Ramus et al., 2018; Williams et al., 2020).

An alternative approach to comparing clinical and healthy groups would be to compare clinical groups with population norms, as is done with well-established cognitive dimensions, such as general intelligence, personality, and psychopathology scales (Beck et al., 1996; Costa Jr. & McCrae, 2008; Wechsler et al., 2008). If neuroanatomical norms for a population were available, then comparing any clinical group to these norms would overcome the issue of sampling variability in the control group. However, neuroanatomical norms are not easy to establish, as they require large populations, and neuroanatomical measures often depend on MRI scanner characteristics and acquisition sequences. Valid norms would therefore only be established within a given study in a single scanner, or in a small set of comparable scanning sites with similar acquisition protocols as done by the UK Biobank.

For this reason, the UK Biobank, the largest neuroimaging dataset available to date (N ∼ 40 000), is an ideal candidate to create neuroanatomical norms that could be re-used for multiple studies of neurological and psychiatric disorders. These norms should be sex-specific, given that the two sexes differ on a number of neuroanatomical brain measures (Kaczkurkin et al., 2019; Lotze et al., 2019; Ritchie et al., 2018; Ruigrok et al., 2014; Sanchis-Segura et al., 2019), and have different risks of developing certain neurological and psychiatric disorders (Beam et al., 2018; Boyd et al., 2015; Seedat et al., 2009). And age should also be considered, as it is associated with variations in neuroanatomical measures (Fjell et al., 2013; Hurtz et al., 2014; Narvacan et al., 2017; Vinke et al., 2018; Wierenga et al., 2014), cognitive function (Simon R. Cox et al., 2018; S.R. Cox et al., 2019), and disease risk (Fiske et al., 2009; Fjell et al., 2009; Jellinger & Attems, 2015).

Finally, global brain measures should be taken into account to create norms that are independent of variations in brain size. Although there is mounting evidence that brain allometry - the non-linear scaling relationship between regional and global brain dimensions - is an inherent property of the brain (Finlay et al., 2001; Jäncke et al., 2015a; Liu et al., 2014; Mankiw et al., 2017; Reardon et al., 2018; Toro et al., 2009; Williams et al., 2020), standard modes of adjustment for individual differences in global measures, such as the proportion method, or linear covariate adjustment, omit brain allometry. To this day, numerous studies have shown that different methods of adjustment for brain size contribute to the variability of reported volumetric group differences (Lefebvre et al., 2015; O’Brien et al., 2006, 2011; Reardon et al., 2016; Sanchis-Segura et al., 2019) and some specifically suggest that omitting brain allometry leads to spurious group differences (Mankiw et al., 2017; Reardon et al., 2016; Williams et al., 2020). Since regional/global relationships follow a power function in a majority of regions, it is recommended to log-transform regional and Total Cerebral Measures (TCMs; i.e., Total Brain Volume (TBV), Total Mean Cortical Thickness (MCT), or Total Surface Area (TSA)) to account for allometric scaling and obtain a more accurate description of the relationship between brain regions and TCMs.

Thus, the present study’s first aim is to produced neuroanatomical norms in the UK Biobank that take into account sex, age, and the allometric relationships between regional and global brain measures. Our second goal is to investigate the extent to which neuroanatomical variations depend on sex, age (linear and quadratic), and brain allometry effects and their interactions. Finally, our third aim is to compare TCM adjustment techniques to examine whether omitting brain allometry systematically biases reported results. By generating neuroanatomical markers across volumes, mean thicknesses, and surface areas available in the UK Biobank, the present paper provides UK population norms for future studies that aim to link regional neuroanatomical markers to specific cognitive and behavioral traits or neurological and psychiatric disorders.

## 2. Methods

### 2.1. Participants

Participants were drawn from the UK Biobank, an open-access large prospective study with phenotypic, genotypic, and neuroimaging data from 500 000 participants recruited between 2006 and 2011 at 40 to 69 years old in Great Britain (Sudlow et al., 2015). All participants provided informed consent (“Resources tab” at https://biobank.ctsu.ox.ac.uk/crystal/field.cgi?id=200). The UK Biobank received ethical approval from the Research Ethics Committee (reference 11/NW/0382) and the present study was conducted based on application 46 007.

Currently, Magnetic Resonance Imaging (MRI) data and Imaging-Derived Phenotypes (IDPs) are available for about 41 000 participants. This study analyzed the IDPs from the first imaging visit generated by an image-processing pipeline developed and run by the UK Biobank Imaging team (Alfaro-Almagro et al., 2018; Miller et al., 2016).

#### 2.1.1. Brain Image Acquisition and Processing

A standard Siemens Skyra 3T running VD13A SP4 with a standard Siemens 32-channel RF receive head coil was used to collect data (<u>Brain Scan Protocol</u>). The 3D MPRAGE T1-weighted volumes were analyzed by the UK Biobank Imaging team with <u>pipeline scripts</u> that primarily call for FSL and Freesurfer tools. Details of the acquisition protocols, image processing pipeline, image data files and derived measures IDPs of brain structure and function are available in the <u>UK Biobank Imaging Protocols.</u>

#### 2.1.2. Total Brain Volume (TBV)

TBV was calculated as the sum of the total grey matter volume (GMV; i.e., sum of cortical and subcortical GMV, data-field 26518), cerebellum white matter volume (WMV, data-fields 26556 for left and 26587 for right), and cerebral WMV (data-fields 26553 for left and 26584 for right) from the UK Biobank ASEG Freesurfer segmentations. Refer to Supplemental Info 1 for more on the choice of TBV. Individuals with missing data for these regions were excluded from the analyses, yielding 40 055 participants.

#### 2.1.3. Scanner Site

The age and sex of participants differed across the 3 scanner sites located in Cheadle (Site 11025), Reading (Site 11026), and Newcastle (Site 11027; See Supplemental Info 1). One individual without scanner site was removed from the analyses yielding 40 054 participants.

#### 2.1.4. Sex

Participants who did not self-report as male or female or whose self-reported sex and genetic sex differed were also excluded from the analyses (N = 26). When genetic sex was not available, reported-sex was used to define the sex of the participant. Of the 40 029 participants included in the analyses, there were more females (N= 21 142) than males (N = 18 886, χ^2^(1) = 127.15, p < 2.2e-16).

#### 2.1.5. Age

To obtain a continuous and more precise measure of age, age was calculated based on the year and month of birth of the participant and the day, month, and year of their MRI visit. Mean age was at 63.70 years old (*SD* = 7.54). Males (*M* = 64.39 years, *SD* = 7.65) were older than females (t _(39180)_ = -17.18, p < 2.2e-16, *M* = 63.09 years, *SD* =7.39).

### 2.2. Image Derived-Phenotypes (IDPs)

The descriptive statistics of all global and regional IDPs analyzed in the present study and their respective data-fields and segmentation origin are listed in Supplemental Table A1. The majority of IDPs correspond to grey matter, since white matter volumes were not segmented by the UK Biobank Imaging team.

#### 2.1.1. Global IDPs

A total of 9 global IDPs were investigated: TBV, Total Mean Cortical Thickness (Total MCT), Total Surface Area (TSA), Subcortical GMV, Cortical GMV, Cerebral WMV, Cerebellar GMV, Cerebellar WMV, and the Brainstem volume.

WMV measures were obtained by summing left and right global measures from Freesurfer ASEG segmentations (data-field <u>190</u>). Cerebellum GMV was calculated as the sum of the cerebellar volumes from the FAST segmentations (data-field <u>1101</u>). Total MCT and TSA were respectively calculated as the sum of the mean cortical thickness and surface area measures from the Freesurfer a2009s segmentations (data-field <u>197</u>). The whole brain stem measure was taken from the Freesurfer subsegmentations (data-field <u>191</u>) and the Subcortical GMV measure was calculated as the sum of the left and right whole amygdala, hippocampus, and thalamus volumes from the Freesurfer subsegmentations (data-field <u>191</u>) and the left and right caudate, accumbens, pallidum, and putamen of the Freesurfer ASEG segmentations (data-field <u>190</u>).

Based on the recommendations from the <u>UK Biobank Imaging Protocols</u>, we excluded Freesurfer IDPs when T2-FLAIR was not used in addition to the T1 images to obtain the segmentations from Freesurfer a2009s (volume, surface area, and mean thickness) and Freesurfer subsegmentations. Moreover, 790 individuals had missing values for all FAST cerebellum segmentations and were excluded from the FAST segmentation analyses.

Thus, while 40 028 individuals were included in the analyses for TBV, Cerebellum WM, and Cerebral WM, 39 238 individuals were included in the analyses with FAST segmentations and 38 710 were included in the analyses of the Freesurfer Subsegmentations and Freesurfer a2009s segmentations. Missing values and null segmentations (e.g., 0 mm3) for a region were replaced by the mean of that region when calculating global measures. See Supplemental Info 3 for correlations between the global measures provided by Freesurfer ASEG and those calculated from the FAST, Freesurfer Subsegmentations, Freesurfer ASEG, and Freesurfer a2009s Segmentations.

#### 2.2.2. Regional IDPs

A case-wise participant exclusion strategy was applied to each IDP for the regional analyses: participants with a missing value or a segmentation error for a region were excluded from the analyses of that region but were maintained in the analyses of other IDP. Following visual examination of the distribution of regional cerebral measures, values 3 times the inter-quartile range for a region were considered to be segmentation errors and were removed from the analyses of that region.

A total of 620 regional IDPs were investigated: 444 cortical regions (148 volumes, 148 surface areas, and 148 cortical thicknesses) from the Freesurfer a2009s segmentations (Destrieux Atlas, data-field <u>197</u>), 116 whole segmentations and subsegmentations of the amygdala, hippocampus, and thalamus and subsegmentations of the brainstem (Freesurfer subsegmentations, data-field <u>191</u>), 28 cerebellum GMV segmentations from the FAST segmentations (data-field <u>1101</u>), and 32 subcortical, white matter, and ventricle volumes from the Freesurfer ASEG segmentations (data-field <u>190</u>). Freesurfer subcortical segmentations for the caudate, putamen, accumbens, and pallidum were used instead of the preregistered FIRST volumes, for segmentation consistency with the other subcortical and cortical volume which were segmented from Freesurfer.

### 2.3. Statistical Analyses

Analyses were preregistered on OSF and run using R (R Core Team, 2019). The preregistration and code are on OSF (https://osf.io/s4qc5/?view_only=bb067d96d0df4ae4902f99747d60e828). Used packages are listed in Supplemental Info 7.

#### 2.3.1. Data Transformation

To examine allometric scaling, regional and global measures were log10 transformed. All continuous variables were centered around the mean in order to examine the effects of a variable when other variables are at their mean value. The categorical sex variable was coded -0.5 for males and 05 for females. The scanner site variable was dummy coded with the largest site, Cheadle (Site 1102), as reference.

#### 2.3.2. Analyses

Global and Regional analyses were performed twice. Once without scaling (dividing by 1 SD) to obtain the allometric scaling coefficient for each brain region and once with scaling to report standardized betas as effect sizes. Non-linear effects of age were modeled with quadratic age over spline regression as spline regressions do not yield interpretable beta coefficients of age. Isometry was tested using the linear hypothesis function which tests if scaling coefficients obtained without scaling variables differ from 1. Scanner site (Cheadle - Site 1102, Reading - Site 11026, and Newcastle - Site 11027) was additionally added as a covariate, although it was omitted from the preregistration.

#### 2.3.3. Global Analyses

Global analyses were conducted to evaluate how TBV varies with age, age^2^, and sex (equation 1) and how TSA, Total MCT, Cerebral GMV, Cerebral WMV, Total Subcortical GMV, Cerebellum GMV, Cerebellum WMV, and the brainstem volume vary with TBV, age, age^2^, and sex (equation 2).

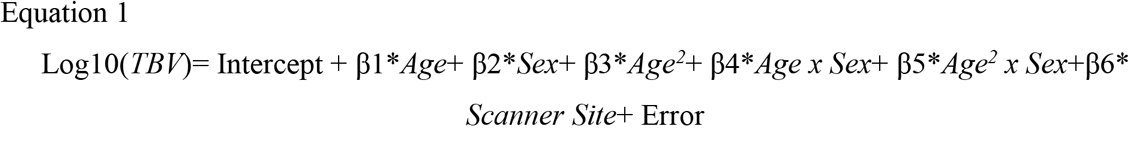

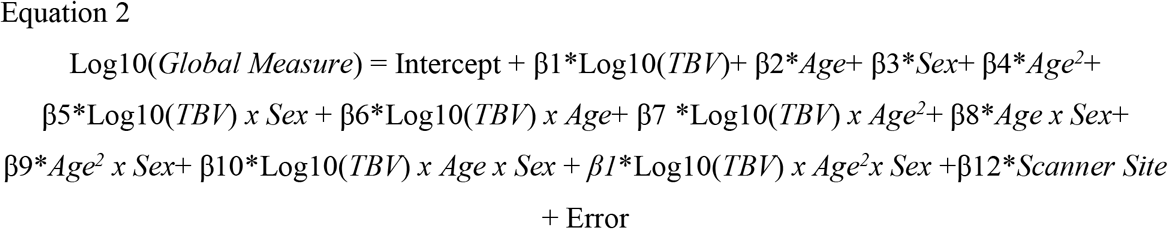

#### 2.3.4. Regional Analyses

Regional analyses were conducted to evaluate how regional volumes, surface areas, and cortical thicknesses vary with TCM, age, age^2^, and sex with equation 3. The TCMs were TBV for volumes, total MCT for mean thicknesses, and TSA for surface areas.

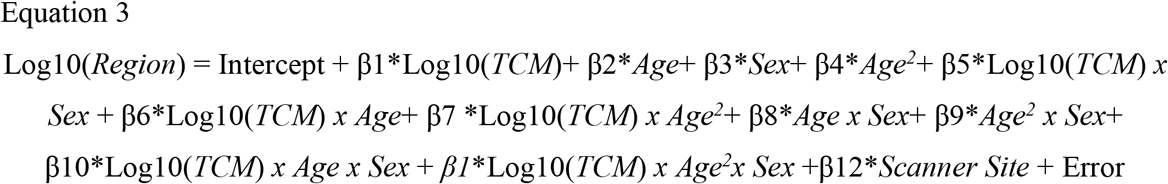

#### 2.3.5. Person-level Neuroanatomical Markers

To obtain a global and regional marker of an individual’s deviance from the norm in terms of volume, mean thickness, and surface area, we extracted the residuals from each dependent variable. The residuals were obtained from the model where continuous variables were centered but not scaled to maintain differences in magnitude across regions. An individual’s residual value for a given regional measure reflects that individual’s deviance from the norm, given his/her age, sex, and TCM, and therefore constitutes a new neuroanatomical marker.

#### 2.3.6. Person-level Global Neuroanatomical Deviance

From the person-level local neuroanatomical markers, we generated four person-level global neuroanatomical deviance measures with equation 4: one for volumes, one for mean thicknesses, one for surface areas, and one for all regions. The person-level global neuroanatomical deviance measure corresponds to a person’s global neuroanatomical deviance from the norm. Although we pre-registered equation 4 without dividing by the total number of investigated regions for a global measure (N), we did so to obtain a value reflecting mean deviation relative to the norm. Considering that all IDPs were not available for all individuals, we excluded participants with more than 10% of missing data across regional IDPs. The cerebral marker across brain measures, which was not preregistered, was calculated by averaging the Z-score of the volumetric, cortical mean thickness, and cortical surface area global neuroanatomical deviance marker.

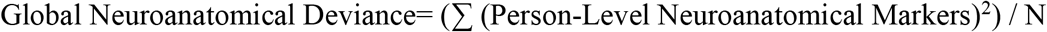

#### 2.3.7. Exploratory Analyses

These analyses were not preregistered unless otherwise stated (for more details see Supplemental Info 4). In brief, we first examined whether global and regional measures were allometric with the linearHypothesis function from the car R package (Fox et al., 2020) with the null hypothesis being “The slope of log10(TCM) is equal to 1”. Then, we examined whether the scaling coefficient of mean thicknesses with TBV differed from ⅓ and whether the scaling coefficient of surface areas with TBV differed from ⅔ with the linearHypothesis function (Fox et al., 2020) and appropriate null (e.g., null hypothesis for mean thicknesses: “The slope of log10(TBV) is equal to 1/3”). We would expect these coefficients if brain growth was proportional (similar to a sphere) and if larger brains were scaled-up versions of smaller brains. Third, we examined sex differences in variance with a Levene’s test (F-test) and calculated as the variance ratio as Female SD / Male SD. Fourth, we compared the results of our main analysis to those obtained when using a linear covariate TCM adjustment (i.e., equation 4 without the log transformation) and when using the proportion adjustment for TCM (i.e., dividing a region by TCM to obtain an adjusted region measure and running equation 4 on the adjusted region without the log transformation and the main effect of TBV). Fifth, as preregistered, we attempted to replicate previous studies on cerebral sex differences that considered brain allometry. Sixth, we replicated Ritchie and colleagues’ (2018) analyses of the Desikan-Killiany cortical measures and FIRST subcortical volumes with the linear covariate TCM adjustment. We additionally ran the same analyses with the allometric TCM adjustment to examine whether we observed the same effects of omitting brain allometry when investigating sex differences in the UK Biobank using different cerebral segmentations and statistical analyses.

#### 2.3.8. Multiple Comparison Corrections

Considering that 620 regions and 11 beta coefficients were investigated for regional IDPs, three thresholds of significance were used: 0.05/11, 0.05/620, and 0.05/ (11 * 620). The same rationale was applied to the global IDPs, with the following thresholds for TBV 0.05/5, 0.05/9, and 0.05/ (5 * 9) and for the remaining global measures: 0.05/11, 0.05/9, and 0.05/ (11 * 9). Significant variance ratio differences are reported at p < 0.05/629 (sum of global and regional cerebral measures).

## 3. Results and Discussion

To avoid redundancy, results are sequentially discussed. Only results at the strictest level of significance are reported in text: p < 0.05/ (5*9) for TBV, 0.05/ (11*9) for other global measures, p < 0.05/ (11*620) for regional measures, and p < 0.05/629 for sex differences in variance. Scaling coefficients (α) correspond to the estimate when continuous variables are centered, and standardized betas (β) reflect the estimate when continuous variables are centered and scaled (1 SD).

Descriptive statistics are available in Supplemental Tables A2-14. Regression results by region are available in Supplemental Tables B1-B10 and by main effectsor interaction, in Supplemental Tables C1-C15. See Supplemental Tables D2-D5 for statistics on regional deviance from isometry. Correlations between the left and right region in terms of the scaling coefficient with the TCM, the age standardized betas, and sex standardized betas are available on OSF, in Supplemental Figures File 1(https://osf.io/s4qc5/?view_only=bb067d96d0df4ae4902f99747d60e828) and correlations between cortical scaling, sex, and age coefficients are available in Supplemental Table C1.

### 3.1 Allometry

#### 3.1.1 Global Allometry

All global scaling coefficients were hypoallometric (α ranging from 0.03 to 0.91), suggesting that these regions increase less than TBV as TBV increases, except for cerebral WMV, which was hyperallometric (α = 1.21). The scaling coefficient of TSA with TBV significantly differed from the theoretical value 2/3 (α = 0.89) and the scaling coefficient of Total MCT with TBV differed from 1/3 (α = 0.03). In cerebellar GMV, all regions were hypoallometric (α = 0.50 - 0.95), except for 4 isometric regions (α = 0.93 - 0.95). Ventricles and the cerebral spinal fluid were hypoallometric (α = 0.40 - 1.01), except for the lateral ventricles (left α = 1.01 and right α = 0.96). The optic chiasm, corpus callosum, cerebellum WMV, and ventral diencephalon measures were hypoallometric (α = 0.55 - 0.93), whereas for the mid-anterior segmentation of the corpus callosum which was isometric (α = 0.99).

Allometric coefficients were generally consistent with previous studies that report hyperallometry in cerebral WMV (Jong et al., 2017; Toro et al., 2009) and hypoallometry in the majority of cerebellar (Mankiw et al., 2017), subcortical (Jong et al., 2017; Liu et al., 2014; Reardon et al., 2016; Williams et al., 2020), and corpus callosum volumes (Lefebvre et al., 2015). See Supplemental Table D1 for global deviance from allometry.

#### 3.1.2 Cortical Allometry

TBV was a significant positive predictor of all volumes, TSA of all surface areas, and Total MCT of all mean thicknesses. Scaling coefficients varied across regions and measures (Table 1). The scaling coefficients of cortical volumes were highly correlated to those of cortical surface areas (*r* = 0.85, p < 2.2e-16) but were not correlated to those of cortical mean thicknesses (*r* = - 0.05, p = 0.535). In cortical regions, 98 volumes, 62 areas, and 63 mean thicknesses were hypoallometric (α = 0.60 - 0.96), while 18 volumes, 50 areas, and 58 mean thicknesses were hyperallometric (α = 1.04-1.62), and 32 volumes, 36 areas, and 27 mean thicknesses were isometric (α = 0.94-1.11; Figure 1).

**Table 1.**
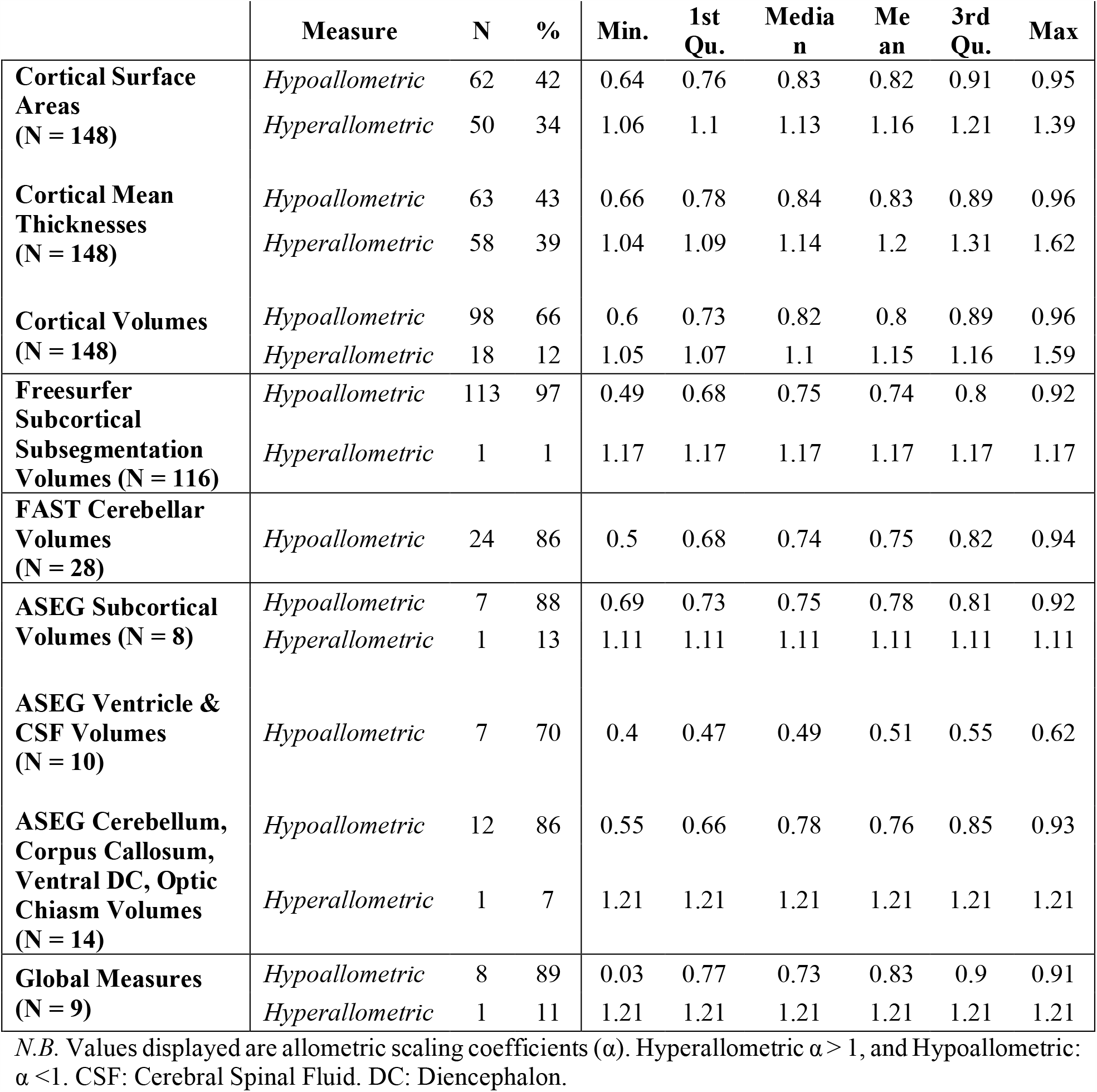
Cerebral Regions Exhibiting Brain Allometry.

**Figure 1.**
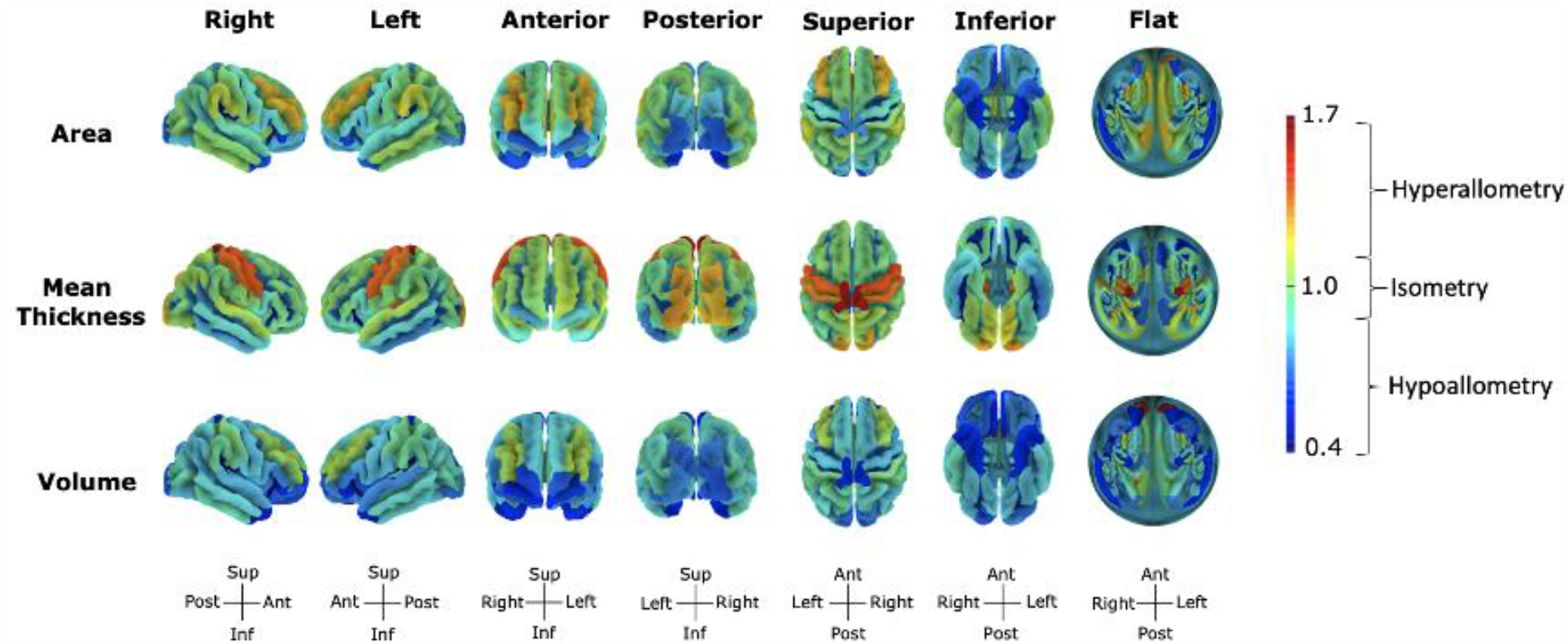
Scaling Coefficients of Cortical Surface Areas, Mean Thicknesses, and Volumes with Total Brain Volume (TBV). Values are the scaling coefficients of a region with TBV and range from 0.61 (volume of the right posterior ramus of lateral sulcus) to 1.63 (mean thickness of the left paracentral gyrus and sulcus). The flat representation corresponds to the flattened image of the superior view with the midline of the circle reflecting regions within the sagittal plane and circle edges reflecting inferior regions. Figures made with https://neuroanatomy.github.io/cortex/ (Toro, 2020).

**Figure 2.**
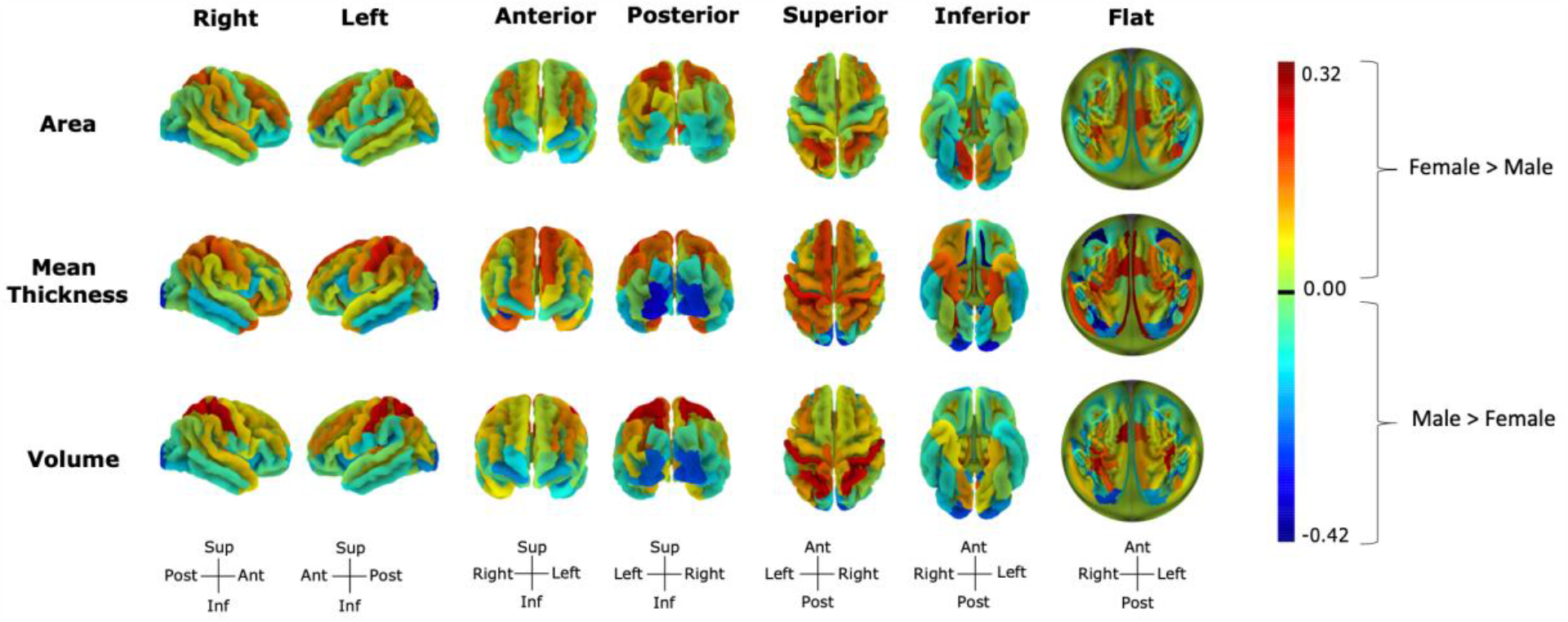
Sex Differences across Cortical Measures. Sex effects differences from -0.40 (mean thickness of the left medial orbital sulcus) to 0.32 (mean thickness of the left transverse temporal sulcus). Negative effects reflect greater male than female volumes. The flat representation corresponds to the flattened image of the superior view with the midline of the circle reflecting regions within the sagittal plane and circle edges reflecting inferior regions. Figures made with https://neuroanatomy.github.io/cortex/ (Toro, 2020).

These findings mirror those reported by Liu and colleagues (2014) and Reardon and colleagues (2018), who found that scaling across cortical regions is heterogeneous, covering a wide range of hypoallometry, isometry, and hyperallometry values. Similarly to Reardon and colleagues (2018), who studied the scaling relationship between vertex area and cortical surface area, surface areas were hyperallometric in the middle frontal and supramarginal gyri and sulci, and hypoallometric in the sensorimotor cortices (precentral, paracentral, and postcentral gyri and sulci), occipital temporal regions, and some cingulate (anterior, mid-anterior, and post dorsal) and callosal (sub and pericallosal) regions.

#### 3.1.3 Subcortical Allometry

All subcortical volumes were hypoallometric (α = 0.49 - 0.92) except for the right lateral posterior and the right limitans suprageniculate nuclei, which were isometric (α = 0.95, α = 1.08, respectively), and the left limitans suprageniculate nuclei and left accumbens area, which were hyperallometric (α = 1.17, α = 1.11, respectively).

General hypoallometry across whole subcortical structures is consistent with previous findings (Jäncke et al., 2015b; Liu et al., 2014; Williams et al., 2020). As for allometry within subcortical structures, Reardon and colleagues (2018) are the only ones to date that examined and reported variations in scaling within subcortical structures. However, the latter study examined the scaling relationship of the vertex area of subcortical subregions with cortical area, whereas the present study examined volumetric variations within these regions. Thus, our finding that allometry varies within subcortical volumes extends our understanding of the cerebral scaling relationships between regional and global measures.

#### 3.1.4 Mean Thickness and Surface Area Scaling with TBV

Exploratory analyses on the scaling relationship between TBV and cortical mean thicknesses or surface areas revealed that the scaling coefficients of all mean thicknesses with TBV were different from one-third, whereas the scaling coefficients of 19 regional surface areas with TBV did not differ from two-thirds (Supplemental Tables D6-7).

This study adds the literature suggesting that larger brains are not simply a scaled-up version of smaller brains (Finlay et al., 2001; Im et al., 2008; Toro et al., 2008), as the first study to examine scaling coefficients of regional cortical surface areas and mean thicknesses with TBV that go beyond lobar segmentations. If the brain regions grew proportionally to brain size, total MCT would scale to the power of one-third with TBV, and TSA to the power of two-thirds with TBV (Finlay et al., 2001). Yet, we find that total MCT scaled to the power of 0.03 with TBV and that the majority of cortical mean thicknesses had allometric scaling coefficients close to 0, reflecting the stability of cortical mean thicknesses with TBV growth. Moreover, TSA scaled to the power of 0.89 with TBV and the majority of cortical surface areas had hypoallometric scaling coefficients that were greater than two-thirds. The greater than geometrically expected hypoallometry across surfaces can be explained by the dramatic increase in gyrification (Fish et al., 2017), and more specifically, in sulcal convolution, that occurs with the expansion of TBV (Im et al., 2008; Toro et al., 2008). Finally, the heterogeneity of allometric patterns observed across the cortex may be explained by the nonuniform gyrification of the cortical surface (Fish et al., 2017).

#### 3.1.5 Sex or Age-Dependent Allometry

Allometry depended on sex (0.2%, 12/628) or age (11%, 71/628) in relatively few regions (details in Supplemental Info 5.1). Sex differences in allometry were less frequent, although larger (|α| = 0.05 - 0.12), than age differences in allometry (|α| = 0.01 - 0.04). Regional volumes generally increased less with TBV in females, while mean thicknesses increased less with Total MCT in males. Although sparse, the presence of TCM interactions with age or sex suggest that matching individuals between groups by TCM may not be appropriate for all regions, as cerebral sex differences may reflect sex-dependent distributions of tissues instead of individual differences in brain size (Luders et al., 2009). Considering that these interactions are often overlooked in studies examining sex and age differences (e.g., Ritchie et al., 2018; Vinke et al., 2018), we suggest that they be considered to obtain unbiased estimates of age and sex effects on the brain and to accurately identify associations between brain regions and behavioral or cognitive traits.

#### 3.1.6 Conclusion on Allometry

All brain regions varied with global brain measures, and the majority (86%) were allometric as they scaled non-linearly with their TCM. Of the regions exhibiting allometry, hypoallometry was reported in 93% of volumes, 55% of the cortical surface areas, and 52% of the cortical mean thicknesses (Table 1). While the association between scaling and cognition and behavior remains unknown, our study adds to the literature (Finlay et al., 2001; Jäncke et al., 2015b, 2019; Jong et al., 2017; Toro et al., 2009) supporting allometric scaling as an inherent property of the brain that varies across regions and cerebral measures, and provides scaling coefficients for regions that were not previously investigated (e.g., subcortical subsegmentations, ventricles etc.).

### 3.2 Sex Differences

#### 3.2.1 Global Measures

TBV was significantly larger in males than in females (β = - 1.14). Once TBV was adjusted for with the allometric adjustment (equation 3), the cerebellar WMV (β = 0.27), cerebellar GMV (β = 0.25), total MCT (β = 0.12), and cerebral WMV (β = 0.06) were greater in females, while brainstem volume (β = -0.21), total subcortical volumes (β = -0.08), TSA (β = -0.07), and cerebral GMV (β = -0.02) were greater in males.

Consistent with previous studies, males had a larger TBV (e.g., Ritchie et al., 2018; Ruigrok et al., 2014), while females had a relatively larger Total MCT (Im et al., 2008, p. 200; van Velsen et al., 2013). However, we did not find that males and females had a similar TSA (Im et al., 2008), nor did we observe greater cerebral WMV in males relative to brain size (Chen et al., 2007; Gur et al., 1999). Instead, our analyses revealed that males have a larger TSA and females a greater cerebral WMV. Considering the small magnitude of these sex differences (<0.1) and the sample size of the previous studies (N < 150), we speculate that these studies were not sufficiently powered to reliably estimate these effects.

#### 3.2.1 Cerebellar GMV

The Freesurfer ASEG Cerebellum GMV – used to calculate TBV - was larger in males (β = - 0.36), whereas the FAST Cerebellum GMV - calculated as the sum of the cerebellar lobes and vermes from FAST Diedrichsen Cerebellar Atlas - was larger in females (β = 0.25).

Result discrepancies between segmentation algorithms may stem from Diedrichsen’s segmentation algorithm ignoring individual WMV and GMV intensities, which are taken into account by Freesurfer. Or differences may be due to Freesurfer over-labelling peripheral tissue as it is more sensitive in regions of low contrast between tissue types (Carass et al., 2018). Although the cerebellar GMV of the FAST and ASEG segmentations only correlated at 76%, discrepancies across cerebellar segmentations did not influence our measure of TBV, as the TBV calculated by summing regional segmentations (including the FAST GMV) correlated at 99.8% with our measure of TBV (derived from the ASEG segmentations).

#### 3.2.3 Regional Cerebellar GMVs

When examining sex differences in the cerebellum with the FAST cerebellum regional segmentations, females had larger cerebellar GMV in 82% of regions (23/28) with coefficients ranging from 0.08 (Right Crus I) to 0.64 (Vermis X). The crus I vermis and the left and right lobule V did not differ between sexes and the left and right lobule X were larger in males (β = -0.17, β = -0.12, respectively).

Our findings contrast with the literature on sex differences within cerebellum (review in Han et al., 2020). For instance, Han and colleagues (2020) instead reported that the VIIIA lobules were larger in males and that the right I–III lobules, and IX and X Vermes were larger in females when adjusting for intracranial volume with the linear covariate method. Moreover, in our replication of Mankiw and colleagues’ (2017) study, we found that, instead of being greater in males, the cerebellum, flocculus, cerebellar lobule VIIb, VIIb, and VIIA, and Crus II volumes were greater in females and that the flocculus volume did not vary between sexes. Although both studies used segmentation algorithms that have a better parcellation accuracy than the SUIT segmentation (Diedrichsen et al., 2009) of the present study (Carass et al., 2018; S. Han, Carass, et al., 2020), discrepancies in the literature may also stem from differences in sample age (mean age = 12.5 and 70 years old, respectively) and size (N = 116 and 2,023, respectively).

In light of the difficulty of segmenting the cerebellum and the differences in regional specificity and accuracy across segmentations, we suggest that future studies take advantage of the large UK Biobank dataset to apply and compare cerebellar segmentation algorithms.

#### 3.2.4 Whole Subcortical Volumes

The thalamus (Right β = -0.15, Left β = -0.08), putamen (Left and Right β = -0.18), left pallidum (β = -0.08), and left amygdala (β = -0.12) were larger in males and the hippocampus (Right β = 0.07, Left β = 0.06) and left accumbens were larger in females (β = 0.10). The right pallidum volume, the caudate volumes, and the right accumbens area volume did not differ between sexes.

Our findings are consistent with previous studies reporting greater thalamic volume in males (Lotze et al., 2019) and larger hippocampal volumes in females (Malykhin et al., 2017; Nordenskjöld et al., 2015), although they contrast with research supporting the absence of sex differences in the amygdala (Lotze et al., 2019) and the hippocampus (Ritchie et al., 2018; Tan et al., 2016) or greater male hippocampal volume (Lotze et al., 2019; Pintzka et al., 2015). And yet, we similarly find that males have larger putamen, pallidum, and left amygdala volumes in our replication of studies examining sex differences when considering brain allometry (Reardon et al., 2016; Sanchis-Segura et al., 2019).

We additionally replicated Ritchie and colleagues’ (2018) findings that males had greater pallidum, putamen, and amygdala volumes, when adjusting for TBV with the linear covariate adjustment and analyzing FIRST subcortical segmentations. For a detailed analysis of the replication by region see Supplemental Info 6.4.2 and Supplemental Tables G5-9.

When adjusting for TBV with the linear covariate or the allometric approach in the replication models with the FIRST segmentations, we found similar subcortical sex differences to those reported in our main analyses with the Freesurfer segmentations with some exceptions. Specifically, instead of being larger in males, the left thalamus was larger in females and right thalamus did not show sex differences in the replication models. Moreover, the right accumbens area was larger in females and the right pallidum was larger in males in the replication analyses, although they did not differ between sexes in the main analyses. These discrepancies may stem from the different terms and interactions included in the main and replication analyses or from differences between the FIRST and Freesurfer subcortical segmentations. For instance, FIRST provides a segmentation of the amygdala more similar to that of manual tracing than Freesurfer (Morey et al., 2009), although the amygdala agreement with manual segmentation is relatively poor compared to other regions such as the hippocampus (Morey et al., 2010), potentially due to the complexity of the structure (Schoemaker et al., 2016).

Sex differences from our main analyses additionally varied across hemispheres. For instance, the amygdala volume was larger for males in the left hemisphere and similar across sexes in the right hemisphere. However, seemingly inconsistent results on whole subcortical structures may be illuminated by examining their subcomponents, as provided by the Freesurfer subcortical subsegmentations.

#### 3.2.5 Subcortical Subsegmentations

Sex differences were found in 67% of the subcortical subsegmentations (74/110), with greater male volume in 42 (38%) regions and greater female volume in 32 (29%) regions (details in Supplemental Info 5.2.3).

The magnitude of the subsegmentation subcortical sex differences was not perceptible at the whole subcortical level due to the presence of sex differences in opposite directions. For instance, medial and lateral regions of the thalamus were considerably larger in females (β ranging from 0.06 to 0.25), although the whole thalamus volume was greater in males (Right β = -0.15, Left β = -0.08). Moreover, the right cortical nucleus (β = 0.13) was larger in females even though sex differences were absent in the whole right segmentation of amygdala. Taken together, these findings support a high variability of sex differences within subcortical structures and highlights the importance of favoring fine-grained segmentations, as done in this study, to better understand where cerebral sex differences lie.

#### 3.2.6 Cortical Regions

Cortical sex differences were present in 57% of surface areas (84/148) and 66% of volumes (97/148) and mean thicknesses (98/148). No clear spatial trend in sex differences was apparent across lobes. In terms of cortical volumes, 62 regions (42%) were larger in males, ranging from - 0.27 (right occipital pole) to -0.06 (right inferior temporal sulcus), and 35 regions (24%) were larger in females, ranging from 0.06 (right precentral gyrus) to 0.26 (right postcentral gyrus). As for cortical mean thicknesses, 50 regions (34%) were greater in males and 48 (32%) were greater in females. Greater mean thicknesses in males varied from -0.39 (left medial orbital – olfactory-sulcus) to -0.05 (parieto-occipital sulcus or fissure), while greater mean thicknesses in females ranged from 0.06 (left transverse frontopolar gyri and sulci) to 0.31 (left transverse temporal sulcus). Finally, in terms of cortical surface areas, males had larger surface areas in 50 regions (34%) ranging from -0.22 (right medial occipito-temporal-collateral – sulcus and lingual sulcus) to -0.06 (left precentral gyrus), and females had larger surface areas in 34 regions (23%) with coefficients ranging from 0.06 (left superior frontal sulcus) to 0.21 (right anterior transverse temporal gyrus of Heschl; Figure 3).

**Figure 3.**
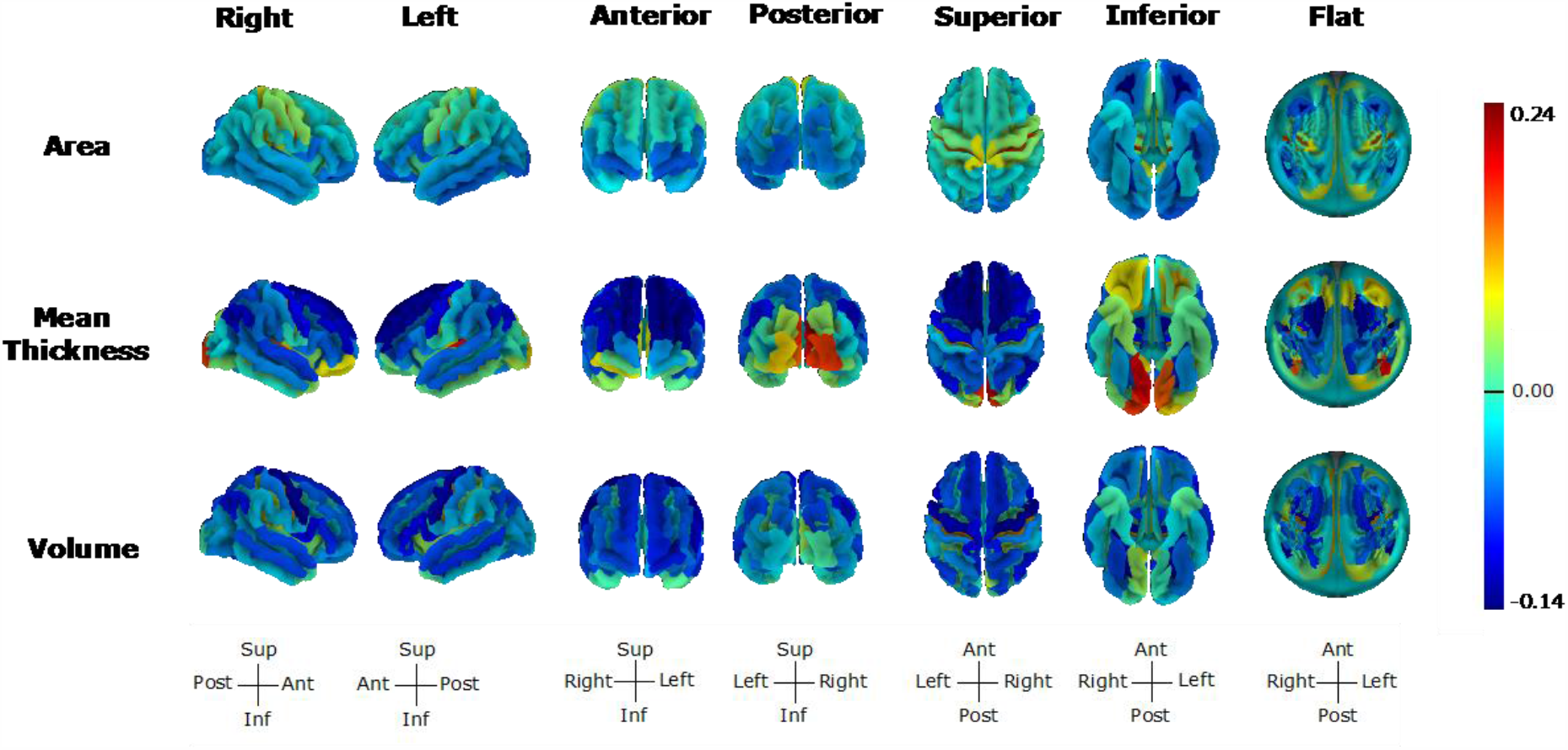
Linear Age Effects across Cortical Measures. Age effects ranged from -0.40 (mean thickness of the right inferior segment of the circular sulcus of the insula) to 0.22 (mean thickness of the right cuneus gyrus, O6). The flat representation corresponds to the flattened image of the superior view with the midline of the circle reflecting regions within the sagittal plane and circle edges reflecting inferior regions. Figures made with https://neuroanatomy.github.io/cortex/ (Toro, 2020).

We replicated the majority of sex differences (88%, 90/98) reported by Ritchie and colleagues (2018) with the Desikan-Killiany Cortical segmentations when adjusting for TCM with the linear covariate approach. We observed significant sex differences in additional regions (81%, 150/186), possibly due to our larger sample size. Sex differences from the main analyses with the Destrieux segmentation and the replication analyses with the Desikan-Killiany appeared to be generally consistent.

In line with a study of cortical volumetric sex differences in 411 middle aged participants (Chen et al., 2007), we found that males had a larger left inferior temporal gyrus and larger right occipital lingual and right middle temporal gyri, while females had a larger right inferior parietal gyrus, right post-dorsal part of the cingulate gyrus and sulcus, and left and right mid-anterior and post-ventral parts of the cingulate gyrus. Consistent with a study of 2 838 middle aged adults (Lotze et al., 2019), we found that females have larger volumes in the superior parietal lobe and right orbitofrontal cortex, whereas males have larger volumes in the left temporal pole and right fusiform gyrus. Yet, in contrast with the previous literature on sex differences in GMV (Lotze et al., 2019; Ritchie et al., 2018; Ruigrok et al., 2014), we reported that females have a larger precentral gyri than males and that males have a larger right anterior part of the cingulate gyrus and sulcus than females. Moreover, we did not find sex differences in the left anterior part of the cingulate gyrus and sulcus, although past studies suggest that this volume was larger in females (Lotze et al., 2019; Ruigrok et al., 2014). The divergence in some of the presented results can partly be attributed to differences in sample size, varying neuroimaging techniques, and to the small size of these effects (median |β| = 0.09) as well as to differences in sample age range. And yet, we observed a similar pattern of sex differences in our replication of Ritchie and colleagues’ (2018) analyses with the allometric adjustment for TCM, suggesting that the different terms included in our models and the differences between the Destrieux and the Desikan-Killiany Atlases had little influence on the majority reported cortical sex differences.

#### 3.2.7 Sex Differences in Variance

In addition to observing mean sex differences in two-thirds of regions, we found sex differences in variance in 49% (306/629) of regions. A total of 253 (40%) regions exhibited greater male variability and 56 (9%) exhibited greater female variability (Table 2). Sex differences in variance ranged from 0.82 (for the right cerebellar lobule VIIIa, implying greater male variability) to 1.17 (for the optic chiasm; Supplemental Info 6.1 and Supplemental Tables E1-7).

**Table 2.** Variance Ratios of Sex Differences across Segmentations

| *Segmentation* |  | **Sex Differences in Variance Ratio** |  | **min** | **1st Qu.** | **median** | **mean** | **3rd Qu.** | **max** |
| --- | --- | --- | --- | --- | --- | --- | --- | --- | --- |
|  |  | **N** | **%** |  |  |  |  |  |  |
| ***Cortical Volumes***<br>***(N = 148)*** | *M>F* | 63 | 43 | 0.91 | 0.94 | 0.95 | 0.95 | 0.96 | 0.97 |
|  | *M<F* | 5 | 3 | 1.02 | 1.04 | 1.05 | 1.07 | 1.08 | 1.14 |
| ***Cortical Surface Areas***<br>***(N = 148)*** | *M>F* | 71 | 48 | 0.90 | 0.94 | 0.95 | 0.95 | 0.96 | 0.97 |
|  | *M<F* | 8 | 5 | 1.03 | 1.04 | 1.04 | 1.06 | 1.08 | 1.14 |
| ***Cortical Mean Thicknesses***<br>***(N = 148)*** | *M>F* | 26 | 18 | 0.9 | 0.95 | 0.95 | 0.95 | 0.96 | 0.97 |
|  | *M<F* | 36 | 24 | 1.03 | 1.05 | 1.06 | 1.06 | 1.07 | 1.15 |
| ***Cerebellar Volumes***<br>***(N = 28)*** | *M>F* | 23 | 16 | 0.82 | 0.90 | 0.94 | 0.92 | 0.95 | 0.97 |
|  | *M<F* | 2 | 1 | 1.04 | 1.04 | 1.04 | 1.04 | 1.04 | 1.04 |
| ***Subcortical Subsegmentations***<br>***(N = 116)*** | *M>F* | 64 | 55 | 0.88 | 0.93 | 0.94 | 0.94 | 0.96 | 0.97 |
| ***ASEG Subcortical Volumes***<br>***(N = 8)*** | *M>F* | 2 | 25 | 0.95 | 0.96 | 0.96 | 0.96 | 0.96 | 0.96 |
| ***ASEG Ventricle & CSF Volumes***<br>***(N = 10)*** | *M>F* | 2 | 20 | 0.95 | 0.95 | 0.95 | 0.95 | 0.95 | 0.96 |
|  | *M<F* | 2 | 20 | 1.07 | 1.08 | 1.09 | 1.09 | 1.1 | 1.11 |
| ***ASEG Cerebellum, Corpus Callosum, Ventral DC, Optic Chiasm Volumes***<br>***(N = 14)*** | *M>F* | 5 | 36 | 0.93 | 0.94 | 0.95 | 0.95 | 0.95 | 0.96 |
|  | *M<F* | 3 | 21 | 1.02 | 1.03 | 1.04 | 1.08 | 1.1 | 1.17 |
| ***Global Measures***<br>***(N = 9)*** | *M>F* | 1 | 11 | 0.90 | 0.90 | 0.90 | 0.90 | 0.90 | 0.90 |

*N.B.* Values represent Variance Ratios. M: Male, F: Female. Variance Ratio = Female SD / Male SD. Qu.: Quartile. CSF: Cerebral Spinal Fluid.

As reported by previous studies (Ritchie et al., 2018; Wierenga et al., n.d., 2018, 2019), the majority of brain regions with sex differences in variance were more variable in males (82%) compared to females (18%). Mean thicknesses were generally more variable in females, while volumes and surface areas were more variable in males. Overall, cerebellar lobes and vermes were also more variable in males. Sex differences in variance across subcortical subsegmentations were greater in males in the hippocampus, amygdala, and thalamus. In terms of whole subcortical volumes, both the hippocampus, which was larger in females, and caudate, which did not show mean sex differences, were more variable in males. Finally, regions that were larger in one sex were typically more variable in the other sex.

Differences in variance thus do not appear to be a mechanical consequence of differences in mean, and instead may reflect a distinct phenomenon known as the greater male variability hypothesis. This hypothesis states that males are more variable than females across a variety of psychological and physical characteristics (Ellis, 1894) and is widely supported by a range of human (e.g., Johnson et al., 2008; Ju et al., 2015; Karwowski et al., 2016; Lehre et al., 2009; Wierenga et al., 2019) and animal (e.g., Branch et al., 2020; DeCasien et al., 2020) studies. Although the mechanisms behind the greater male variability hypothesis exceed the scope of the present study, our findings further support greater male variability, which extends well beyond brain measures.

#### 3.2.8 Conclusion on Sex Differences

Overall, we found that sex differences in the brain are the rule rather than the exception, affecting two-thirds (419/629) of the investigated brain measures, with 231 regions relatively greater in males. The standardized coefficients (β) of the sex effect of cortical volumes were highly correlated to those of cortical surface areas (*r* = 0.77, p = 2.28e-43) and moderately so for cortical mean thicknesses (*r* = 0.45, p = 1.23e-08).

Although many of the regional sex differences had very small effect sizes (β < 0.1) and were only significant due to the large sample size, 46% of these cerebral measures (292/629) had a sex difference above 0.1. Specifically, sex differences in cerebellar GMV and WMV, Total MCT, the corpus callosum, and the ventricles were generally greater than 0.1 (details in Supplemental Info 5.2), whereas sex differences in cerebral GMV and WMV, TSA, 51% of subcortical regions and 35 - 45% of the cortical regions were under 0.1.

### 3.3 Age Effects

#### 3.3.1 Global Measures

All cerebral measures decreased linearly with age (β ranging from -0.04 in to -0.33), except for the brainstem volume, TSA, and cerebral WMV which increased (relatively to TBV) with age (β = 0.13, β = 0.11, β = 0.04, respectively). The quadratic age term did not significantly predict cerebral GMV and WMV or the brainstem volume. However, total subcortical volume (β = 0.01) and TSA (β = 0.02) positively, and TBV (β = -0.05), total MCT (β = -0.04), cerebellar GMV (β = -0.05), and cerebellar WMV (β = -0.02) negatively varied with quadratic age.

In line with the literature, TBV and Total MCT decreased with linear and quadratic age (Ritchie et al., 2018; van Velsen et al., 2013; Vinke et al., 2018) and Total MCT decreased more rapidly with linear age in males (van Velsen et al., 2013). Yet, our finding that TSA increased with linear and quadratic age contrasts with previous reports of a decrease in surface area across the lifespan (Hogstrom et al., 2013; Lemaitre et al., 2012; Long et al., 2012). Divergent results between our study and those of Hogstrom and colleagues (2013) and Long and colleagues (2012) can be explained by their omission of brain size, as we similarly observed a decrease of TSA with age when excluding TBV from our models. However, when applying the proportion TCM adjustment, as done by Lemaitre and colleagues (2012), we observed an increase in TSA with age, suggesting that differences between our studies may instead stem from differences in segmentation or sample characteristics (e.g., smaller sample (N = 216), wider age range (18 to 87 years old)). Based on our findings, the relative expansion of TSA in older adults with age may reflect a global spread of the sulci, which appears to occur more rapidly in males than females.

#### 3.3.2 Cerebellar Volumes

There was a linear decline with age of the cerebellar GMV for the FAST (β = -0.20) and the Freesurfer ASEG (β = -0.02) segmentations. However, the FAST cerebellar GMV was also negatively predicted by quadratic age (β = -0.05) and its linear decline with age was quicker in males compared to females (β = 0.03). As for the regional FAST cerebellar GMVs, 27 out of 28 (96%) cerebellar volumes decreased linearly with age. The age effect in the cerebellar IX vermis (β = -0.02) did not reach significance. Linear age effects ranged from -0.20 (right Crus I) to -0.05 (vermis Crus I). We found a negative quadratic age effect in 74% (20/27) of regions with a linear age effect, which ranged from -0.07 (Left Cerebellar Lobule VIIb) to -0.03 (Right Cerebellar Lobule VIIb). The cerebellar IX vermis was the only area with a quadratic age but no linear age effect (β = -0.04).

The linear decrease with age of the cerebellum mirrors previous findings (for review Bernard and Seidler, 2014). The literature also similarly reports a linear rather than a non-linear cerebellar change with age (for review Fjell et al., 2013; Fjell & Walhovd, 2010) and the absence of an age by sex interaction in cerebellar volumes (Hoogendam et al., 2012; Raz et al., 2005) when examining the Freesurfer segmentation of the cerebellum. The discrepancies in results between cerebellar segmentations further highlight the nonnegligible impact that the type of cerebellar segmentation algorithm has on reported results (as discussed in section 3.2.2 and 3.2.3).

Although our findings add to the scarce literature on age related changes within the cerebellum, age effects in these GMVs remain highly inconsistent (Bernard & Seidler, 2013; S. Han, An, et al., 2020; Koppelmans et al., 2017). We speculate that these differences in reported results can be attributed to the insufficient sample size of previous studies to investigate the these effects (our median |β|= 0.17; N = 54 for Bernard and Seidler (2013) and N = 213, for Koppelmans et al. 2017) and differences in segmentation algorithms, which vary in accuracy and in the number of segmented cerebellar regions (Carass et al., 2018; L. Han et al., 2019).

#### 3.3.3 Whole Subcortical and Subcortical Subsegmentation Volumes

The putamen (β = -0.09 for both), accumbens area (Left β = -0.33, Right β = -0.24), amygdala (Left β = -0.16, Right β = -0.14), and hippocampus (Left β = -0.23, Right β = -0.19) decreased with age. The pallidum (Left β = 0.03, Right β = 0.06) and caudate (Left β = 0.12, Right β = 0.15) volumes increased with linear age, whereas the thalamus did not show significant linear age effects. The accumbens area, amygdala, and hippocampus volumes had a negative quadratic age effect, ranging from -0.03 (right accumbens area) to -0.09 (left presubiculum head), and the thalamus, caudate, and right putamen all showed positive quadratic age effects, ranging from 0.02 (right putamen) to 0.06 (left caudate).

Linear age was a significant predictor of 95% (104/110) of subsegmentations and quadratic age was a significant predictor of 88 (104/110) subsegmentations. Although there were no age effects at the whole subcortical level of the thalamus, we found 24 linear age effects with an absolute effect size greater than 0.1 across thalamic subsegmentations. More positive linear and quadratic age effects were found in ventral and intralaminar thalamic volumes. On the other hand, the majority of the amygdala and the hippocampal subsegmentations (>92%) decreased with age. The direction of the quadratic age effects on subcortical subsegmentations was similar to that of the whole subcortical volumes, as we found negative quadratic age effects across the amygdala and hippocampal subsegmentations and positive quadratic age effects across the thalamic subsegmentations.

The volumetric decline in the amygdala, hippocampus, putamen, and nucleus accumbens with age mirrors previous findings (Hogstrom et al., 2013; Kurth et al., 2017; Sele et al., 2020; Vinke et al., 2018). However, our finding of an increase in pallidum volume with age contrasts with past studies reporting a small decrease with age in this region (Sele et al., 2020; Vinke et al., 2018). Moreover, while we add to the literature reporting an expansion of the caudate with age (Vinke et al., 2018), a small decrease has also been reported (Sele et al., 2020). In terms of the discrepancy in the caudate results, we speculate that Sele and colleagues’ (2020) sample (N = 231) was insufficient to observe such small changes with age (β = 0.05-0.06). As for the discrepancies in the pallidum results, differences may be attributed to the different terms included in the regressions, as Vinke and colleagues (2018) modeled non-linear changes with splines instead of a quadratic age.

#### 3.3.4 Cortical Regions

Cortical regions increased with linear age in 22% of volumes, 35% of mean thicknesses, 24% of surface areas, and declined with linear age in 41% of volumes, 52% of mean thicknesses, 33% of surface areas (Figure 3). Cortical volumes decreased linearly with age in 60 regions, ranging from -0.13 (right precentral gyrus) to -0.02 (right parieto-occipital sulcus), and increased with age in 33 regions, ranging from 0.03 (right superior segment of circular sulcus of the insula) to 0.13 (left central sulcus). Cortical mean thicknesses decreased with age in 77 regions, ranging from -0.14 (right inferior circular sulcus of the insula) to -0.02 (left posterior dorsal cingulate gyrus), and increased with age in 52 regions, ranging from 0.02 (left middle occipital gyrus) to 0.22 (right cuneus gyrus). Surface areas increased linearly with age in 36 regions, ranging from 0.02 (left superior frontal gyrus) to 0.15 (right central sulci), and decreased with age in 49 regions, ranging from -0.11 (right the orbital sulci) to -0.02 (left lateral aspect of the superior temporal gyrus). In contrast, a positive quadratic age effect was reported in 5% of volumes, 6% of mean thicknesses, 3% of surface areas, and a negative quadratic age effect was reported in 1% of volumes, 9% of mean thicknesses, 3% of surface areas. For further details on cortical age effects, see Supplemental Word Document Section 5.3.

Our findings coincide with and extend the literature reporting large age-related changes across the cortical measures (e.g., Fjell et al., 2009; Lotze et al., 2019; Pintzka et al., 2015; Salat et al., 2004; Storsve et al., 2014) The majority of frontal volumes decreased with linear age, while frontal surface areas decreased with age in the orbital gyri and sulci and the inferior frontal gyrus and frontal mean thicknesses decreased with age in the frontal medial and frontal superior regions. Temporal surface areas and volumes generally decreased with linear age, whereas age effects on temporal mean thicknesses were more variable. For instance, we observed a mean thickness thinning in the lateral aspect of the superior temporal gyrus and the middle temporal gyrus and a thickness increase of the planum polare, Heschl’s gyrus, lingual gyrus, and the temporal pole. Occipital regions generally decreased with age in surface areas and volumes, and increased with age in mean thicknesses, while parietal regions mainly decreased with age across cerebral measures.

Our findings additionally shed a light on the inconsistent age-related changes reported in motor, somatosensory, and visual cortices (Hogstrom et al., 2013; for review McGinnis et al., 2011). In terms of motor and somatosensory cortices, we found the largest surface area expansions in the paracentral lobule and sulcus and the precentral and postcentral gyrus (motor and somatosensory cortices) and a reduction of a similar size in these regions with age in terms of mean thicknesses and volumes. As for the visual cortices, the largest mean thickness expansions with age occurred in the cuneus gyrus, the lingual gyrus, occipital pole, the inferior, middle, and superior, occipital gyri. The surface areas of these regions generally decreased with age and their volumes were either unaffected by age or showed a slight increase with age.

#### 3.3.5 Conclusion of Age Effects

We observed a linear change with age in 76% (480/629) and a quadratic one in 25% (159/629) of regional cerebral measures. About 49% of regions decreased with linear age and 28% increased with linear age. Regions showing quadratic age effects typically showed linear age effects. A sex by age interaction was observed for regional measures in 14% of regions (87/629), ranging from -0.16 (right cerebellar lobule X) to 0.19 (left fimbria). For detailed results on the ASEG and Freesurfer subsegmentations linear and quadratic age effects, see Supplemental Info 5.3 and for detailed results and a discussion of the sex by age and sex by age^2^ interactions see Supplemental Info 5.4.

### 3.4. Does Brain Allometry Influence Reported Results?

We examined the effects of omitting brain allometry and adjusting for TCM with different methods. To do so, we compared the number of significant results from our main analyses obtained with the allometric TCM adjustment to those obtained when using the linear covariate or proportion TCM adjustment. The models with the linear covariate TCM adjustment yielded similar significant effects and interactions (i.e., age and sex effects and their interactions with and without TCM) to the allometric models: The linear covariate model under- or overestimated effects in 2.35% (102/4340) of statistical tests and these differences occurred in regions near significance with small effect sizes (Supplemental Figures File 3 and 4, https://osf.io/s4qc5/?view_only=bb067d96d0df4ae4902f99747d60e828). In contrast, the proportion adjustment for TCM overestimated 14.24% (618/4340) of effects and interactions reported in our main analyses compared to the allometric TCM adjustment (see Supplemental Info 6.2.1 and Supplemental Table F1 for details). In our replication of the FIRST subcortical and Desikan-Killiany Cortical sex differences reported by Ritchie and colleagues (2018), the models with the linear covariate and the allometric approach additionally yielded consistent results, except for the sex differences of the left transverse temporal and the left pars opercularis volumes (β =0.02 for both), which only reached significance in the allometric model. Finally, based on our correlational analyses of a region’s deviance from isometry (i.e., |1 - scaling coefficient|) and the difference in the effect size of a term between models with varying TCM adjustments (e.g., |Proportion Sex β - Allometric Sex β|), we found that discrepancies in significance between the linear and allometric models were accentuated in more allometric regions, specifically for the proportion models (see Supplemental Info 6.2.3, Supplemental Tables F5, and Supplemental Figures File 5 https://osf.io/s4qc5/?view_only=bb067d96d0df4ae4902f99747d60e828).

Sex differences in variance differed in 38.06% (236/620) of regions between the allometric and the linear covariate approach and in 29.03% (180/620) of regions between the allometric and the proportion approach. These discrepancies additionally lead to a change in the direction of the correlation between a region’s sex effect standardized beta and variance ratio. When adjusting for TCM with the proportion or the linear covariate approach, regions that were larger in males became more variable in males, instead of being more variable in females (see Supplemental Info 6.2.2 and Supplemental Tables F2-4). We found a similar change in the direction of the correlations between the sex effect Cohen’s d and variance ratios in our replication of Ritchie and colleagues’ (2018) study with the cortical Deskian-Killiany and subcortical FIRST segmentations (Supplemental Table G10).

In line with previous research, we find more consistent results between the linear covariate and allometric approach compared to the proportion and allometric approach (Mankiw et al., 2017; Reardon et al., 2016; Sanchis-Segura et al., 2019). Our findings additionally suggest that the major source of variation in mean results across models with differing TCM adjustments is due to the omission of the intercept of the relationship between a region and its TCM (as in the proportion method) rather than the omission of its non-linear relationship (as in the proportion and linear covariate methods). Moreover, as the first study to examine the effects of omitting brain allometry on sex differences in variance, we find that omitting brain allometry leads to over or underestimating sex differences in variance depending on the region. Therefore, we suggest that brain allometry generally be considered to provide unbiased estimates of age, sex, and TBV effects and interactions across all brain regions, and stress that previous reported sex differences in variance relative to brain size be reexamined with brain allometry (Ritchie et al., 2018; Wierenga et al., n.d., 2018, 2019).

### 3.5 Neuroanatomical Norms

Neuroanatomical norms were generated for 40 028 UK Biobank participants at two levels: for global brain measures (total cerebral and cerebellar GM and WM volumes, TSA, Total MCT, and total subcortical and brainstem volumes) relative to TBV, and for each regional measure relative to its corresponding TCM (i.e., TBV, TSA, or Total MCT). These norms correspond to the residuals from the full statistical models and reflect the extent to which the cerebral measures of an individual deviates from other individuals of the same sex, age, and total brain size. In addition to creating global and regional neuroanatomical norms for each of the 629 brain measures, we computed global neuroanatomical deviance markers, which reflect the deviance of an individual from the norm across all brain regions. This was done separately for volumes, surfaces and thicknesses, as well as across the three types of measures.

An individual with a “normal” brain - where all regional measures are at the expected value given this individual’s sex, age, and total cerebral size - should have a global neuroanatomical deviance of 0 for volume, surface and thickness. However, values ranged from 0.57 to 1.53 for volumes, 0.02 to 0.05 for mean thicknesses, and 0.04 to 0.13 for surface areas, suggesting that it is ‘normal’ to deviate from the cerebral norms (Table 3). In turn, individuals with a global neuroanatomical deviance above the mean had brain regions that deviated from the regional norms more than most individuals, while values below the mean represent reduced deviance from the neuroanatomical norm.

**Table 3.**
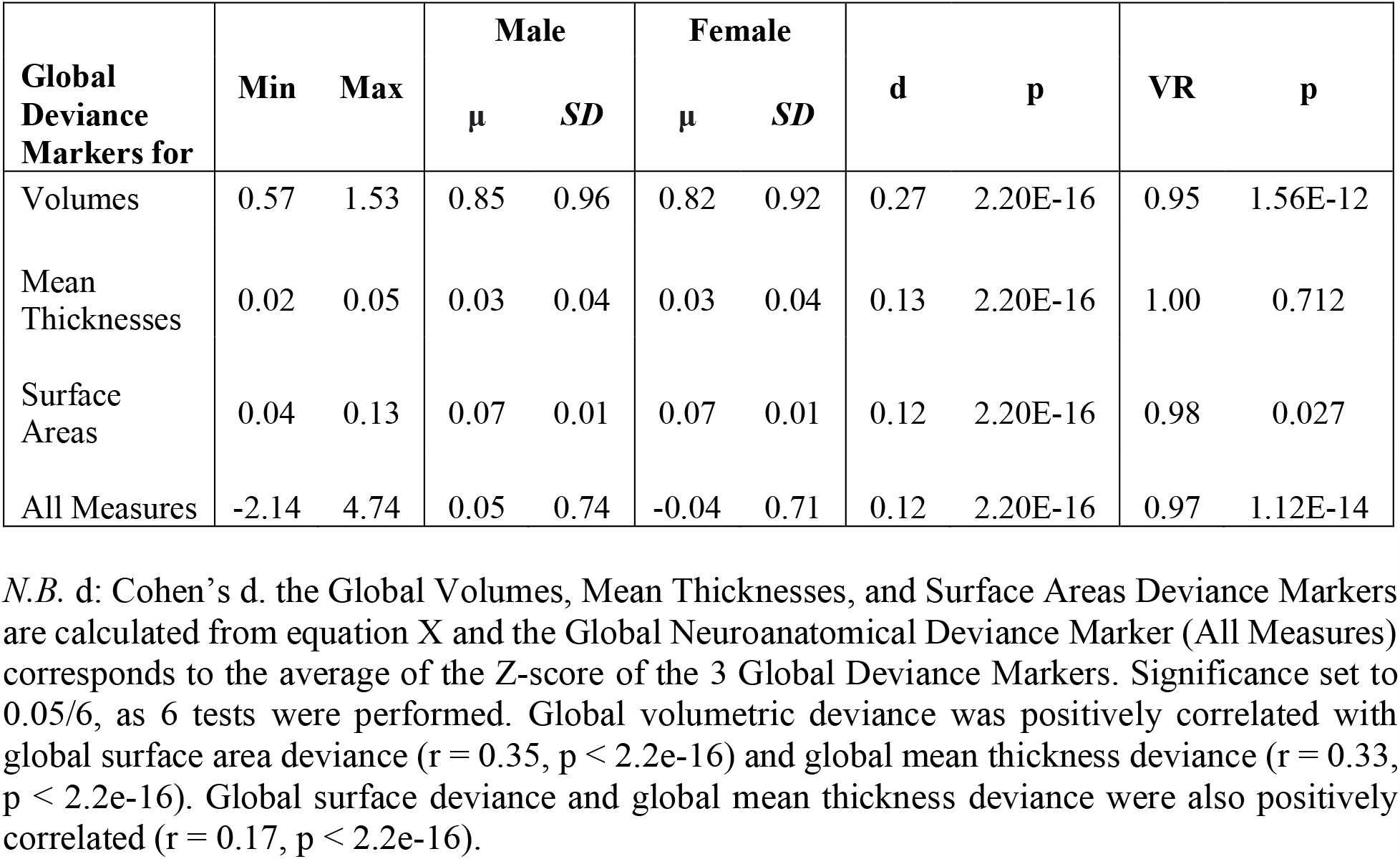
Mean and Variance Sex Differences in across Global Deviance Markers

The means of the volumetric, surface area, and neuroanatomical (all cerebral measures combined) global neuroanatomical deviance were larger in males, suggesting that male volumes and surface areas deviated more from their sex-specific norm than females. However, females deviated more from their norm in the mean thickness global allometry marker. Males additionally had a more variable volumetric and neuroanatomical global allometric markers (Table 3, Figure 4). Thus, investigations of global as well as regional neuroanatomical deviance should always take sex into account.

**Figure 4.**
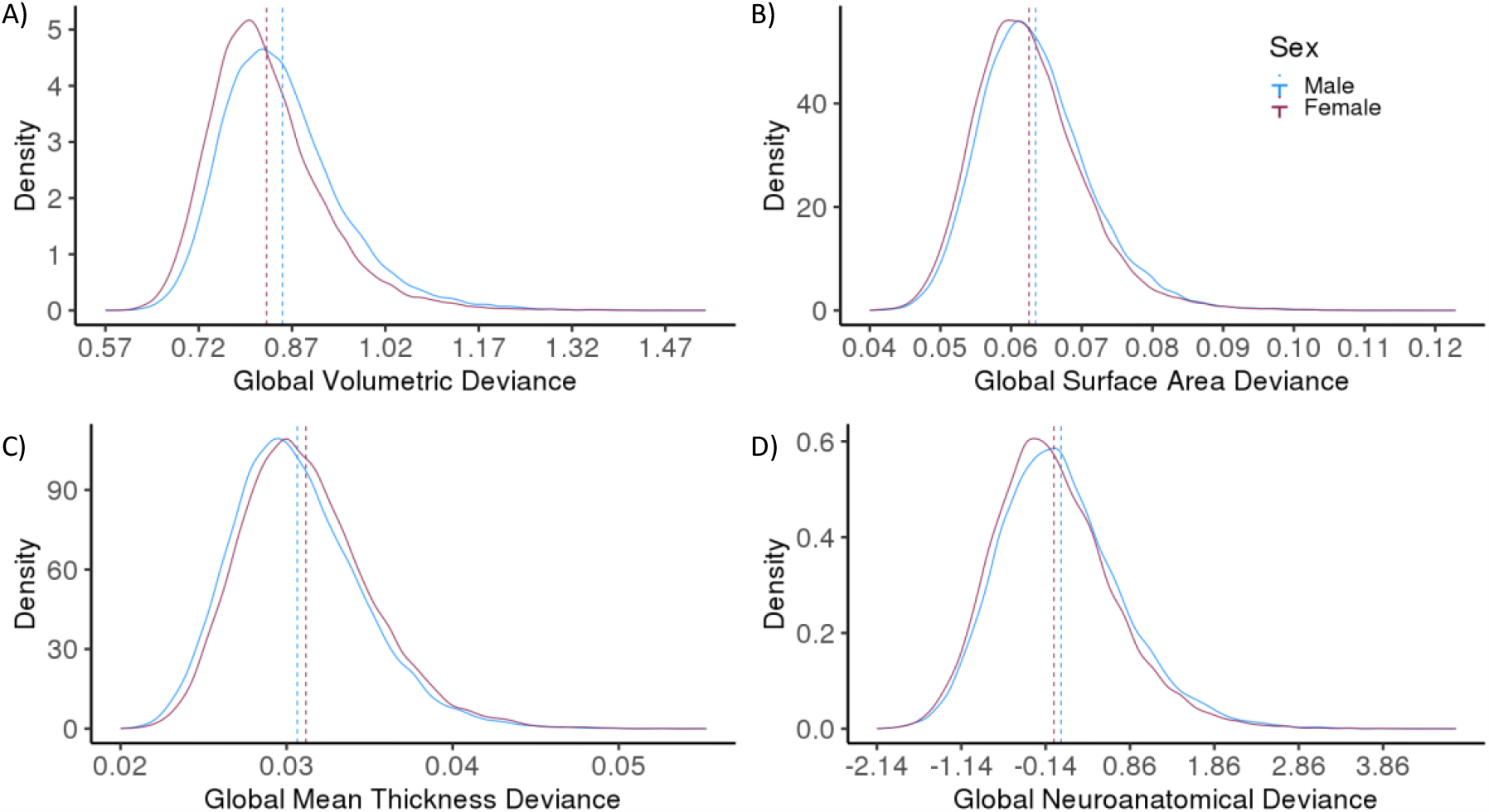
Sex Differences in Global Neuroanatomical Deviances across Volumes (A), Cortical Surface Areas (B), Cortical Mean Thicknesses (C), and all Volumes, Cortical Surface Areas, and Cortical Mean Thicknesses(D). Mean (dashed lines) differences were found across measures, while variance differed between sexes only across volumes. Global Allometry marker corresponds to the square root of the sum of squared residuals divided by the number of regions for that measure from the model with age, total brain volume, and sex as well as age^2^, total brain volume, and sex interactions. Global Neuroanatomical Deviance Marker corresponds to the average of the Z-score of the 3 Global Deviance Markers. Significance set to 0.05/6, as 6 tests were performed.

With the world’s largest neuroimaging dataset, we created neuroanatomical norms in the UK Biobank, to which any UK Biobank individual can be compared, in the same way that scores from intelligence tests or personality questionnaires can be compared to population norms. With these norms, future UK Biobank studies will be able to examine whether individuals that deviate from the norm on a global or regional brain measure also deviate from the norm in terms of cognitive and behavioral traits or of risk for neurological and psychiatric disorders. Having brain markers that are relative to total cerebral measures (rather than raw measures) will make it easier to distinguish the specific contribution of each regional brain measure from that of more global brain measures. As for global neuroanatomical deviance markers, future studies will be able to investigate the extent to which global neuroanatomical deviance reflects disruptions of brain development or serves as a risk factor for neurodevelopmental or psychiatric disorders. For studies examining the associations of specific regional brain measures with cognitive phenotypes, it may be useful to adjust on global neuroanatomical deviance, on top of total brain size, in order to fully dissociate regional from global effects.

### 3.6 Limitations

In light of the “healthy volunteer” selection bias and older age range of the UK Biobank (Fry et al., 2017), the present paper is limited in its capacity to generalize its findings and neuroanatomical norms and markers to other age groups or to the UK population. Moreover, created these norms and markers are not independent of ethnicity or highest level of education attained, factors thought to influence neuroanatomical measures (Shen et al., 2017; Tang et al., 2018). However, this enables future studies to investigate whether the present neuroanatomical markers vary as a function of ethnicity and level of education. If the aim is to generalize these findings to the UK population, we suggest that neuroanatomical markers be created with a representative sample or that weights be used to adjust the phenotypic measures of the UK Biobank to match those of the UK population.

While numerous studies focus on developing machine learning algorithms (e.g., SVM classifications) to generate neuroanatomical markers, we were interested in identifying sex, age, and TCM effects while considering their potential interactions, which are often omitted in the literature. By opting for a regression approach, we were able to quantify effects and interactions for each region, which would have been lost with machine learning. Future studies would nevertheless benefit from examining the age, sex, and global brain effects of other anatomical measures, such as diffusion tractography, and functional measures in the UK Biobank.

While the present study modelled age as a linear and quadratic function, other studies examining sex and age interactions used the nonparametric local smoothing technique (i.e., smoothing splines; Fjell et al., 2013; Vinke et al., 2018), which are thought to be more predictive of individual trajectories and less vulnerable to sampling range (Fjell & Walhovd, 2010). However, this nonparametric approach also requires researchers to make more decisions that contribute to the variability of results. For instance, Fjell and colleagues (2010) initially selected the spline smoothing level that minimized the AIC, but when the absence of smoothing (which is equivalent to the linear least square model) yielded the smallest AIC for the sample of individuals over 60 years old, they chose the smoothing level that minimized the BIC. Considering that the nonparametric local smoothing technique depends on selected parameters and that splines are difficult to interpret if we are interested in quantifying the magnitude of age effects, quadratic age was used instead of splines to model non-linear age in the present study.

## 4. Conclusion

The present study is the largest analysis to date of the age, sex, and TCM effects and interactions on global and regional brain volumes, cortical mean thicknesses, and cortical surface areas. We provide further evidence that brain allometry is a common property of the brain that should be considered to report unbiased estimates of age, sex, and TCM effects and interactions. By generating volumetric and allometric norms in the UK Biobank, we pave the way for future research to examine the associations between these markers and the behavioral and cognitive traits available in the UK Biobank. Once associations between these UK Biobank neuroanatomical norms and cognitive and behavioral measures are established, researchers will be able to examine the extent to which these neuroanatomical markers mediate the effect that genes and the environment have on these traits. This line of research will play critical role in our understanding of the influence that neuroanatomy has on who we are.

## Abbreviations

(TCMs): Total Cerebral Measures
(TBV): Total Brain Volume
(MCT): Total Mean Cortical Thickness
(TSA): or Total Surface Area
(GMV): Grey Matter Volume
(WMV): White Matter Volume

## Acknowledgements

This work received support under the program “Investissements d’Avenir” launched by the French Government and implemented by ANR with the references ANR-17-EURE-0017 and ANR-10-IDEX-0001-02 PSL. Funding was also obtained from Fondation pour l’Audition (FPA RD-2016-8 research grant). This research has been conducted using the UK Biobank Resource. Declarations of interest: none.

## Data/Code Availability

This research has been conducted using data from UK Biobank, a major biomedical database (http://www.ukbiobank.ac.uk/). Restrictions apply to the availability of these data, which were used under license for this study: application 46007. Preregistration and code are available on OSF: https://osf.io/s4qc5/?view_only=bb067d96d0df4ae4902f99747d60e828.

## Conflict of interest

On behalf of all authors, the corresponding author states that there is no conflict of interest.

## Notes

### Competing Interest Statement

The authors have declared no competing interest.

