## Supplemental Information for "Neuroanatomical Norms in the UK Biobank: The Impact of Allometric Scaling, Sex and Age"

### Table of Contents

***1 – Total Brain Volume (TBV) Choice*** ***3***

***2 - Scanner Site*** ***6***

***3 - Correlation between the calculated global measures and those provided by the Freesurfer ASEG Segmentation.*** ***6***

***4 - Exploratory Analyses: Methods.*** ***7***

**4.1. Sex Differences in Variance.** **7**

**4.2. Comparing Results of Different TBV Adjustment Techniques.** **7**

**4.3. Replication of Previous Studies Examining Brain Allometry.** **8**

**4.4. Replication of Ritchie and colleagues (2018) Study and Comparing Results with the Covariate and with the Allometric TBV Adjustment.** **8**

**4.5. Height** **9**

***5 - Supplementary Results and Discussion*** ***10***

**5.1. Age by Total Cerebral Measure (TCM) and Sex by TCM Interactions** **10**

*5.1.1. TCM by Sex.* 10

*5.1.2. TCM by Age.* 12

**5.2. Sex Effects** **13**

5.2.1 *Ventricles and CSF Volumes (ASEG).* 13

5.2.2 *Other Volumes (ASEG).* 13

5.2.3 *Subcortical Freesurfer Subsegmentations.* 13

**5.3 Age Effects** **15**

5.3.1 *ASEG Volumes.* 15

5.3.2 *Subcortical Freesurfer Volumes*. 15

5.3.3 *Cortical Volumes.* 16

5.3.4 *Cortical Mean Thicknesses.* 16

5.3.5 *Cortical Surface Areas.* 16

**5.4 Sex by Age & Sex by Age$^2$** **17**

*5.4.1 Results* 17

*5.4.2 Discussion* 18

***6 - Exploratory Analyses: Results*** ***19***

**6.1 Sex Differences in Variance.** **19**

*6.1.1 Global Measures*. 20

*6.1.2 Cortical Volumes.* 20

*6.1.3 Cortical Mean Thicknesses.* 20

*6.1.4 Cortical Surface Area*. 20

6.1.5 ASEG Volumes. 21

*6.1.6 Subcortical Freesurfer Volumes*. 22

Abbreviations: Total Cerebral Measures (TCMs): Total Brain Volume (TBV), Total Mean Cortical Thickness (MCT), or Total Surface Area (TSA). Grey Matter Volume (GMV). White Matter Volume (WMV).

*6.1.7 Cerebellar GMVs.* 22

**6.2 Comparing TBV Adjustment Techniques – Allometric Versus Covariate or Proportion Adjustment 22**

*6.2.1 Comparing Standardized Beta Coefficients when omitting and considering brain allometry.* 22

*6.2.2 Comparing the Sex Difference in Variance across TCM Adjustment Techniques.* 23

*6.2.3 Correlations between a region's deviance from isometry (i.e., |1 – scaling coefficient|) and the difference in standardized beta between the linear (covariate or proportion) and allometric models* 24

**6.3 Replication of Sex Differences in Studies considering Brain Allometry 25**

**6.4 Replication of Ritchie and Colleagues (2018) Study and Comparing Results with the Covariate and with the Allometric TBV Adjustment. 26**

*6.4.2 Subcortical Volumes: Mean and Variance Sex Differences.* 28

*6.4.3 Cortical Volumes: Mean and Variance Sex Differences.* 28

*6.4.4 Cortical Surface Areas: Mean and Variance Sex Differences.* 28

*6.4.5 Cortical Mean Thicknesses: Mean and Variance Sex Differences.* 29

**6.5 Height 31**

***VI- R packages 31***

### *1 – Total Brain Volume (TBV) Choice*

Both TBV and Total Intracranial Volume (TIV) have been used to adjust for brain allometry to avoid a bias toward isometry when estimating allometric scaling coefficients of large regions (e.g., Jong et al., 2017; Peyre et al., 2020; Reardon et al., 2016; Williams et al., 2020). However, since segmentations yielding the highest number of regions were favored and because we are interested in examining how much a particular volume contributes to the total brain volume at the time of the study (and not at maximal lifetime total brain volume; Herbert et al., 2003; O'Brien et al., 2006), TBV was chosen over TIV. The correlation between TBV and TIV was 88.93%.

We did not use the TBV provided by ASEG (26515) due to a mismatch for some participants between the provided TBV measure (26515) and the one calculated by summing the global measures provided by ASEG: Total GMV Volume (26518) + Cerebellum WMV (Left: 26556, Right: 26587) + Cerebral WMV (Left: 26553, Right: 26584). This mismatch yielded a correlation of 99.73% between TBV measures.

Instead, we used the TBV calculated by summing the global measures provided by ASEG as our measure of TBV since it correlated at 99.82% with the TBV we calculated from summing the regions of interest in the present study.

The ASEG global sum measure was favored over the sum of regions we analyzed in our study to include a maximum number of participants. (We would otherwise, for instance, have to exclude all individuals that do not have the FAST cerebellum segmentations even though they have all the other segmentations).

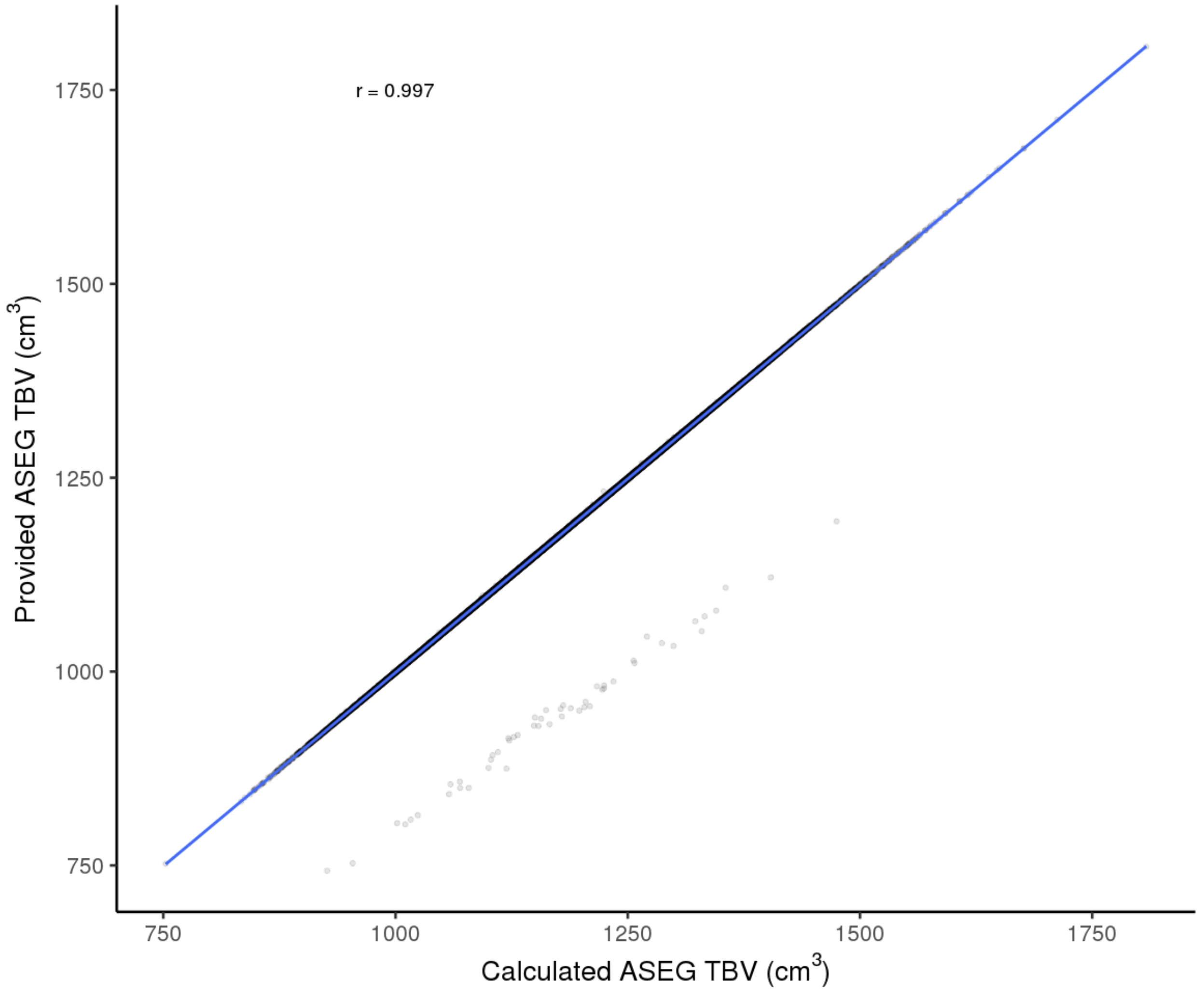


***Figure S1*****. Correlation between the Provided ASEG Total Brain Volume (TBV) Measure and the Calculated Sum of Global Measures from the Freesurfer ASEG Segmentations.** The provided TBV corresponds to data-field 26515 and the Calculated ASED TBV to the sum of Total Grey Matter Volume (data-field 26518), Cerebellum White Matter Volume (Left: data-field 26556, Right: data-field 26587), Cerebral White Matter Volume (Left: data-field 26553, Right: data-field 26584). r, rho = 99.73%.

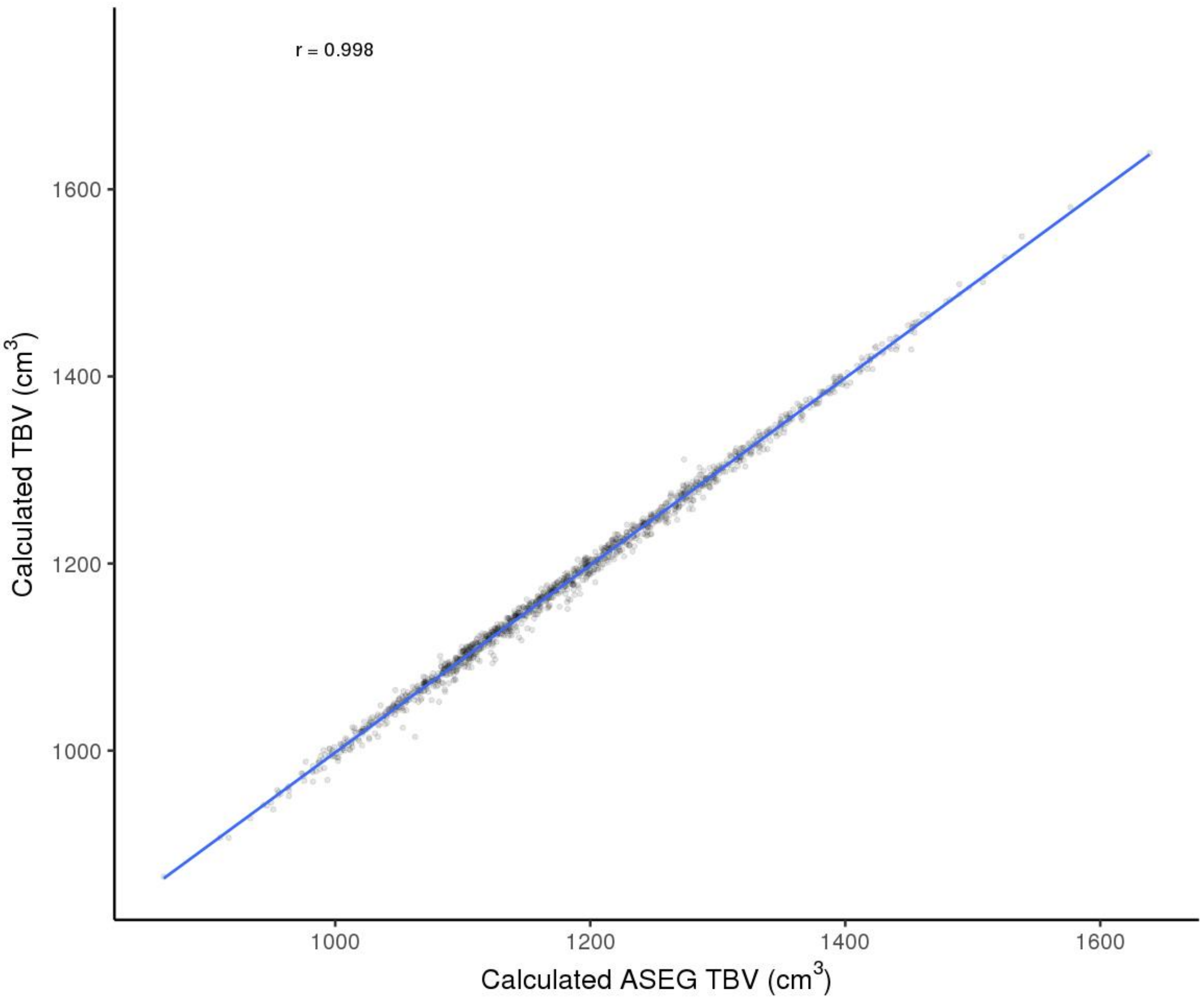


***Figure S2*. Correlation between the Calculated ASEG Total Brain Volume (TBV) Measure and the Calculated TBV from the Regions examined in this study.**

The Calculated ASEG TBV corresponds to the sum of Total Grey Matter Volume (data-field 26518), Cerebellum White Matter Volume (Left: data-field 26556, Right: data-field 26587), Cerebral White Matter Volume (Left: data-field 26553, Right: data-field 26584). r, Pearson r.

#### *2 - Scanner Site*

The number of participants differed across sites (χ2 (2, N = 40 028) = 16 107, $p < 2.2e-16$) and between sexes (χ2 (2, N = 40 028) = 27.1, p =1.314e-06), with more female than male participants across sites (Site 11025: Male N = 12033 and Female N = 12938, Site 11026: Male N = 2318 and Female N = 2752, and Site 11027: Male N = 4535 and Female N = 5452). Age differed between Site 11025 and Site 11026 (b = 2.27 years, *t* (40025) = 19.63, $p < 2.2e-16$), between Site 11025 and Site 11027 (*b* = 1.40 years, *t* (40025) = 15.78, $p < 2.2e-16$), and between Site 11026 and Site 11027 (*b* = -0.87 years, *t* (40025) = -6.70, p = 2.1e-11).

#### *3 - Correlation between the calculated global measures and those provided by the Freesurfer ASEG Segmentation.*

Global measures were calculated as the sum of the regions examined in the present study instead of using predefined measures when available. Since we used regions from different segmentations to perform the most fine-grain analysis possible, we examined the correlation between the summed global measures and the global measures provided by ASEG.

The Total Subcortical (Grey Matter, GM) Volumes measure was calculated as the sum of the amygdala, hippocampus, and thalamus volumes from the Freesurfer subsegmentations (191) and the caudate, accumbens, pallidum, and putamen of the Freesurfer ASEG segmentations (190). This calculated measure correlated at 99.25% with the Total Subcortical Volume measure from the ASEG segmentation (26517).

The Total Cortical GM, which corresponds to the sum of the a2009s Destrieux volumes correlated at 99.97% with the Total Cortical GM provided by the ASEG segmentation.

The calculated total Cerebellum GM, which was obtained by summing the grey matter volumes from the FAST segmentations, correlated with the sum of the left and right cerebellum GM from the ASEG Freesurfer segmentations at 75.83%.

### *4 - Exploratory Analyses: Methods.*

These analyses were not preregistered. Asymmetries in scaling with global measures, and sex and age effects were examined by computing the correlation of the scaling coefficient or the standardized betas of the left and right regions.

#### 4.1. Sex Differences in Variance.

Differences in the variance of cerebral measures between sexes were assessed with a Levene's test (F-test) performed on the person-level neuroanatomical markers. Variance ratios were calculated as the Female SD / Male SD. We report the correlation between the variance ratio and standardized beta of each region to test if the regions showing the largest sex differences also showed the greatest sex differences in variance. We additionally examined differences in the variance ratio of the global allometric marker between sexes.

#### 4.2. Comparing Results of Different TBV Adjustment Techniques.

We investigated whether the significance of the TBV, sex, and age (linear and quadratic) effects and interactions varied with the type of adjustment for TBV. We compared results from the allometric approach to results from the proportion and the covariate approach, separately.

For the proportion method, we obtained an adjusted region measure by dividing a region by its global measure and then applying equation 4 without the main effect of TBV. For the covariate approach, we applied equation 4 to the raw cerebral measures. The main analyses were initially run without controlling for Scanner Site and since the influence of scanner site was limited to a change of 0.01 in beta coefficients of a few regions in the main analyses, Scanner Site was not included in the analyses performed to compare result consistency across three TBV adjustment analyses.

We examined whether the same-sex differences in variance and mean effects and interactions were significant when considering and omitting brain allometry.

To identify if results from allometric models diverged more from those obtained with linear models in regions that were more allometric, we examined the correlation between a region's deviance from isometry (| 1 – scaling coefficient|) and the difference in the standardized beta of an effect and interaction across models with differing TCM adjustments (e.g., |Covariate Sex Std Beta – Allometric Sex Std Beta|). We chose to compare the betas from the effects and interactions that were significant in at least one region in our primary analyses.

#### 4.3. Replication of Previous Studies Examining Brain Allometry.

We attempted to replicate age and sex cerebral differences reported by studies that considered brain allometry. When available we used the same segmentations as these studies for a more accurate replication, although our sample was older and much larger. Specifically, we tried to replicate the following findings:

1. Right and left lobar (frontal, occipital, limbic, parietal, temporal, and cerebellum) sex differences reported by Peyre and colleagues (2020; Figure 1. Right and left cerebellum grey matter: boys greater than girls; right (d = -0.2) and left frontal grey matter: girls greater than boys (d = 0.08-0.1)).

2. A greater pallidal volume in males and relative caudate head expansion and ventral striatum contraction in females were reported by Reardon and colleagues (2016).

3. Relative expansion of total cerebellum, flocculus, and Crus II-lobule VIIIB volumes in males and X-specific contraction of Crus II-lobule VIIIB by Mankiw and colleagues, (2017; Figure 6).

4. Sex differences reported by Sanchis-Segura and colleagues (2019) in Figure 2 when adjusting for allometry with the power-corrected proportion method proposed by Liu and colleagues (2014) were the following:

- Left, female greater than male: Frontal superior, superior orbital, mid, med orbital, occipital med, postcentral, parietal inf, cerebellum X; male greater than female cerebellum crus I, pallidum, putamen, amygdala.
- Right, female greater than male: Frontal superior orbital, frontal med orbital occipital superior, mid, and parietal inferior; male greater than female: temporal pole mid, pallidum, putamen, amygdala, and lingual.

#### 4.4. Replication of Ritchie and colleagues (2018) Study and Comparing Results with the Covariate and with the Allometric TBV Adjustment.

Finally, we attempted to replicate the mean and variance ratio (VR) sex differences reported by Ritchie and colleagues' (2018) while adjusting for age, ethnicity, and linear TBV with the Desikan-Killiany atlas segmentations. We used the same FIRST subcortical volumes segmentations available in the UK Biobank as Ritchie and colleagues (2018). However, we used the Desikan-Killiany Cortical Atlas segmentations from the UK Biobank while Ritchie and colleagues (2018) performed their own Desikan-Killiany Cortical segmentations. To examine if brain allometry influenced reported results with Ritchie and colleagues (2018) model and choice of segmentation, we additionally ran Ritchie and colleagues' (2018) model while adjusting for brain allometry (i.e., log transforming cerebral measures) and compared our results from our replication analyses to those obtained when considering brain allometry. As done by Ritchie and colleagues (2018), we report the correlation between the VR and Cohen's D of each region to examine if mean sex differences show sex differences in variance.

#### 4.5. Height

To assess if global differences were due to differences in height, we also ran equation 2 with log transformed height instead of the log transformed TBV.

Abbreviations: Total Cerebral Measures (TCMs): Total Brain Volume (TBV), Total Mean Cortical Thickness (MCT), or Total Surface Area (TSA). Grey Matter Volume (GMV). White Matter Volume (WMV).

### *5 - Supplementary Results and Discussion*

Since data for the 5th ventricle was only available for 2% of UK Biobank subjects (830/40 028), all 5th ventricle results may not be reliable.

#### 5.1. Age by Total Cerebral Measure (TCM) and Sex by TCM Interactions

##### *5.1.1. TCM by Sex.*

*Global Measures.* Cerebellum GMV increased more with TBV (i.e., showed a larger allometry coefficient α) in males (α = 0.42) compared to females (α = 0.39, β = -0.06).

*Regional Measures.* Only 11 regions out of 620 (0.2%) had significant interactions between total cerebral measure and sex and the interactions were the largest for the following regions. The right fimbria volume increases less with TBV in females (α = 0.29) compared to males (α = 0.50, β = -0.12, SE = 0.02), while the left surface area of the pericallosal sulcus increases more with TSA in females (α = 0.71) compared to males (α = 0.59, β = 0.09, SE = 0.01; Figure S3).

A)

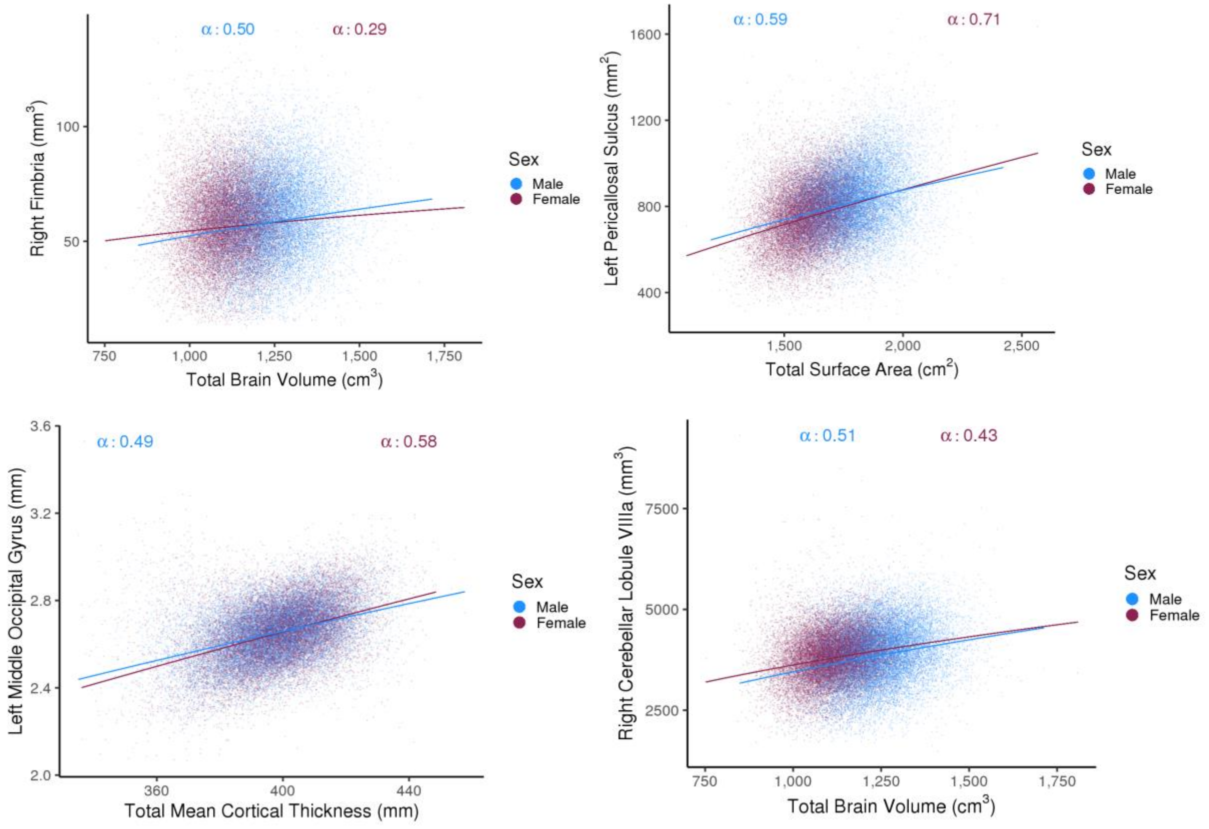

***Figure S3.* Largest Total Cerebral Measure by Sex Interaction**. α is the scaling coefficient of the region with its total cerebral measure. Blue and purple lines correspond to the predicted values of y from the model with TBV, sex, age and TBV, sex, $age^2$ interactions.

***5.1.2. TCM by Age.***

*Global Measures*. Total subcortical volumes and TSA increased more with TBV in individuals 64 years old (median) and under compared to individuals more than 64 years old (β = -0.01 for both). However, Cerebellum GMV and Total MCT increased more with TBV in older adults compared to younger adults (β = 0.03 and β = 0.04, respectively). Total MCT and cerebellar GMV additionally decreased more with age in individuals with a smaller TBV (β = -0.37, β = -0.31, respectively) compared to individuals with a larger TBV.

*Regional Measures.* Only 67 regions out of 620 (11%) had significant interactions between the total cerebral measure and age and were the largest for the following regions. The right fimbria volume increases less with TBV in younger (α = 0.74) compared to older individuals (α = 1.13, β = 0.03, SE = 0.01), while the mean thickness of the left transverse temporal sulcus increases more with the TBV in younger (α = 1.49) compared to older individuals (α = 1.22, β = -0.04, SE = 0.00; Figure S4).

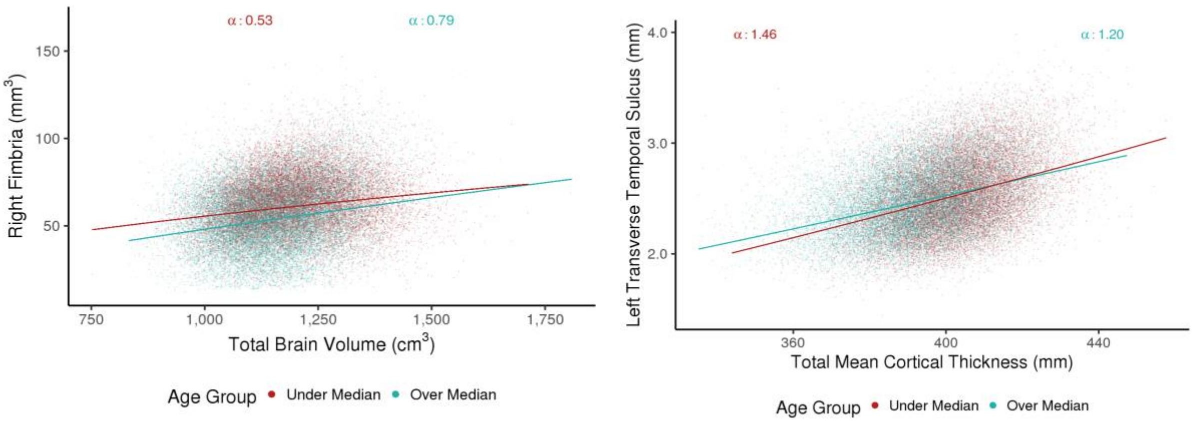


***Figure S4*. Largest Negative and Positive Total Cerebral Measure by Age Effects.** Colored lines correspond to the predicted values of y from the model with TBV, sex, age and TBV, sex, $age^2$ interactions. α is the scaling coefficient of the region with its total cerebral measure.

### 5.2. Sex Effects

***5.2.1 Ventricles and CSF Volumes (ASEG).***

All 9 out of 10 ventricles and CSF segmentations were greater in males ranging from -0.40 (left lateral ventricle) to -0.67 (right choroid plexus), except for the 5$^{th}$ ventricle.

**5.2.2 *Other Volumes (ASEG).***

All 14 remaining ASEG volumes exhibited sex differences. The optic chiasm (β = -0.41), cerebellar GMV (Right β = -0.38, Left β = -0.33), and ventral diencephalon (Right β = -0.15, Left β = -0.15) were larger in males, while cerebellum WMV (Right β = 0.27, Left β = 0.26), cerebral WMV (Right β = 0.05, Left β = 0.07), and corpus callosum (β = 0.27 to 0.40) volumes were larger in females.

**5.2.3 *Subcortical Freesurfer Subsegmentations.***

Of the 80 regions out of 116 subcortical regions (69%) exhibiting sex differences, 46 (40%) were larger in males and 34 (29%) were larger in females. The following subcortical sex differences are described by region.

Sex differences across the brainstem volume subsegmentations ranged from -0.49 (medulla) to -0.10 (pons).

The left amygdala was greater in males than females (β = -0.12), while the right amygdala did not show sex differences. However, sex differences were present in the right amygdala subregions, such as the cortical (β = 0.13) and paralaminar (β = -0.13) nuclei. Overall, 8 amygdala subsegmentations out of 20 were greater in males ranging from -0.18 (left paralaminar nucleus) to -0.06 (left accessory basal nucleus). The 3 regions that were larger in females were the left medial nucleus (β = 0.09) and left and right cortical nuclei (β = 0.09, β = 0.13).

The right and left hippocampus were greater in females than males (Right β = 0.07, Left β = 0.06), although some subsegmentations of the hippocampus were larger in males. Specifically, 5 out of 40 (12.5%) hippocampal volumes were larger in males, including the left and right hippocampal fissure (β = -0.29, β = -0.25), left and right parasubiculum (β = -0.11, β = -0.08), and left CA4 head (β = -0.10). Females showed greater volumes in 17 regions (42.5%) ranging from 0.06 (right subiculum body) to 0.19 (left molecular and granule cell layers of the dentate gyrus).

The right and left thalamus were greater in males (Right β = -0.15, Left β = -0.08), with 42 (81%) regions exhibiting sex differences. Male volumes were larger in 28 out of the 52 thalamic segmentations (54%), ranging from -0.34 (right anterior pulvinar nucleus) to -0.06 (left ventral posterior nucleus), and 14 (27%) regions were greater in females, ranging from 0.07 (left lateral posterior) to 0.25 (right nucleus reuniens of the ventral midline thalamus).

### 5.3 Age Effects

**5.3.1 *ASEG Volumes.***

A total of 28 out of 32 (87.5%) volumes showed linear age effects and 14 out of 32 (43.75%) volumes showed quadratic age effects.

All ventricle and CSF volumes increased with age ranging from 0.18 (4th ventricle volume) to 0.47 (3rd ventricle volume) and were positively associated with $age^2$ ranging from 0.03 (left choroid plexus) to 0.10 (right inferior lateral ventricle). The 5th ventricle did not show linear or quadratic age effects.

There was no linear or quadratic age effect on cerebellum GM, although cerebellum WMV decreased linearly with age (Left $\beta$ = -0.13, Right $\beta$ = -0.11). Volumes of the ventral diencephalon (Left $\beta$ = -0.08, Right $\beta$ = -0.09) additionally decreased with linear age, while cerebral WMV ($\beta$ = 0.04) and the optic chiasm (0.23) increased with linear age.

Interestingly, age effects varied across segments of the corpus callosum. The corpus callosum posterior volume increased linearly with age ($\beta$ = 0.10), while the corpus callosum mid-posterior ($\beta$ = -0.14), central ($\beta$ = -0.20), and mid-anterior ($\beta$ = -0.22) volumes decreased with age and the anterior volume showed no age effect ($\beta$ = 0.00).

Finally, putamen and accumbens area volumes decreased with linear age ($\beta$ = -0.33 to -0.09), while pallidum and caudate volumes increased with linear age ($\beta$ = 0.03 to 0.15). All volumes showed quadratic age effects ranging from -0.04 (left accumbens area) to 0.06 (left caudate), except for the left putamen and pallidum volumes.

**5.3.2 *Subcortical Freesurfer Volumes*.**

Linear age was a significant predictor of 108 out of 116 (93%) subsegmentations and quadratic age was a significant predictor of 94 regions (81%).

The amygdala decreased with age (Left $\beta$ = -0.16, Right $\beta$ = -0.14), with linear age effects ranging from -0.23 (left cortical nucleus) to -0.05 (left paralaminar nucleus), except for the right paralaminar nucleus which did not show linear age effects.

Hippocampal regions decreased with age (Left $\beta$ = -0.23, Right $\beta$ = -0.19), with 37 out of 40 (92.5 %) regions showing linear age effects ranging from -0.29 (right fimbria) to -0.06 (right parasubiculum). In contrast, the left and right hippocampal fissure increased with age (Left $\beta$ = 0.28, Right $\beta$ = 0.34).

The linear age effect on the left and right thalamus segmentations was not significant, although linear age effects were reported in thalamic subsegmentations ranging from -0.30 (right lateral geniculate nucleus) to 0.03 (right pulvinar anterior nucleus) and from 0.03 (right ventral lateral posterior nucleus) to 0.33 (right limitans suprageniculate nucleus).

Of the 108 regions with a linear age effect, 94 had a quadratic age effect ranging from -0.09 (right presubiculum head) to 0.11 (left centromedian nucleus). The left and right lateral posterior nuclei (both, $\beta = 0.06$), the right paralaminar nucleus ($\beta = -0.02$), and right pulvinar inferior ($\beta = 0.07$) showed a quadratic but not a linear age effect.

The amygdala (Left $\beta = -0.04$, Right $\beta = -0.05$) and its subsegmentations as well as the hippocampus (Left $\beta = -0.23$, Right $\beta = -0.20$) and its subsegmentations generally had a negative quadratic age effect. The whole thalamus left and right had a positive quadratic age effect (Left $\beta = 0.07$, Right $\beta = 0.06$), although the right lateral geniculate, right reuniens of the ventral midline thalamus, and right medial mediodorsal nuclei increased with quadratic age.

**5.3.3 *Cortical Volumes.***

Of the 91 regions with a linear age effect, 5 also had a significant positive quadratic age effect. A quadratic age effect in the absence of a linear effect was also reported for the left parahippocampal gyrus ($\beta = -0.03$), the left superior part of the precentral sulcus ($\beta = 0.03$), and the left anterior part of cingulate gyrus and sulcus ($\beta = 0.03$).

**5.3.4 *Cortical Mean Thicknesses.***

Of the 129 regions with an age effect, 8 regions had a positive and 13 had a negative quadratic age effect. The right straight gyrus (gyrus rectus) was the only area with a quadratic age but no linear age effect ($\beta = 0.02$).

**5.3.5 *Cortical Surface Areas.***

Of the 85 regions with an age effect, 4 regions had a positive and 3 had a negative quadratic age effect. The left temporal pole surface area was the only area with a quadratic age but no linear age effect ($\beta = -0.02$).

### 5.4 Sex by Age & Sex by Age$^2$

#### *5.4.1 Results*

*Global Measures.* Total MCT (β = 0.10), cerebellum GMV (β = 0.05) and WMV (β = 0.04) decreased more rapidly with age in males compared to females and TSA increased more rapidly with age in males (β = -0.04; Figure 1A).

*Regional Measures.* A total of 83 regions out of 620 (13%) had a significant sex by age interaction, ranging from -0.16 (right cerebellar lobule X) to 0.19 (left fimbria). Of the 7 regions (1%) with a significant sex by age$^2$ interaction, the choroid plexus volumes (β = 0.08) and right CA4 body (β = 0.05) did not show a significant interaction of age by sex. The largest and only negative sex by age$^2$ effect was reported for the GMV of the cerebellar vermis X (β = -0.10), which also presented an age by sex effect (β = 0.17). The left fimbria and cerebellar vermis volumes decreased more in males than females, while the right cerebellar lobule X decreased more in females than males with age. The left choroid plexus increased more in males than in females with quadratic age (Figure S5).

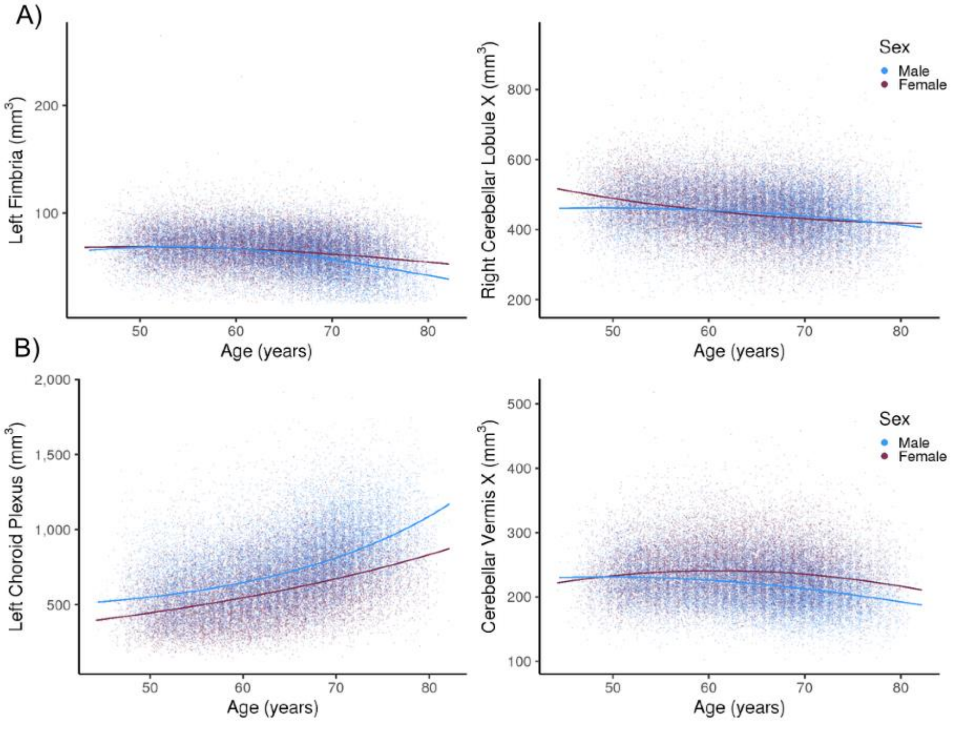


***Figure S5.* Largest Sex by Age (A) and Sex by Age$^2$ (B) Interactions.** Blue and purple lines correspond to the predicted values of y from the model with TBV, sex, age and TBV, sex, age$^2$ interactions.

#### *5.4.2 Discussion*

Sex differences in linear and quadratic age effects were observed for a subset of regions (15%) supporting the presence of sex differences in cerebral aging trajectories (Armstrong et al., 2019; Vinke et al., 2019). Total MCT ($\beta = 0.10$), cerebellum GMV ($\beta = 0.05$) and WMV ($\beta = 0.04$) decreased more rapidly with age in males compared to females and TSA increased more rapidly with age in males ($\beta = -0.04$). Females had a steeper decrease with linear age across occipital temporal mean thicknesses, the thalamus, and its subsegmentations, whereas males showed a more rapid decrease with linear age in frontal and cingulate gyri and sulci mean thicknesses and hippocampal subsegmentations. Females had a steeper decrease with linear age in the cerebellar lobule X, whereas males had a steeper decrease with linear age in the cerebellar lobules iiv, vi, viib, viiia, viiib, ix and vermes viiia, ix, and x. Sex differences in quadratic age effects were only reported in 7 regions including the cerebellum x (flocculus), choroid plexus, and hippocampal CA4 body. Considering that the median absolute effect size for the age by sex interaction was 0.06, we speculate that inconsistent findings with previous studies likely stem from underpowered designs in addition to differences in age modeling (e.g., linear versus spline) and sample characteristics (e.g., age range; Armstrong et al., 2019; Good et al., 2001; Lemaitre et al., 2012; Vinke et al., 2019). Our study further adds to the literature reporting the presence of age-related cerebral changes that vary as a function of sex in late adulthood.

### *6 - Exploratory Analyses: Results*

#### 6.1 Sex Differences in Variance.

Sex differences in variance occurred in 306 out of 629 regions (49%; Supplemental Tables Tables E1-7) across measures types, ranging from 0.82 (for the right cerebellar lobule VIIIa, implying greater male variability) to 1.17 (for the optic chiasm, suggesting greater female variability, Figure S6). A total of 253 (40%) regions exhibited greater male variability and 56 (9%) exhibited greater female variability. A detailed description is made available by segmentation.

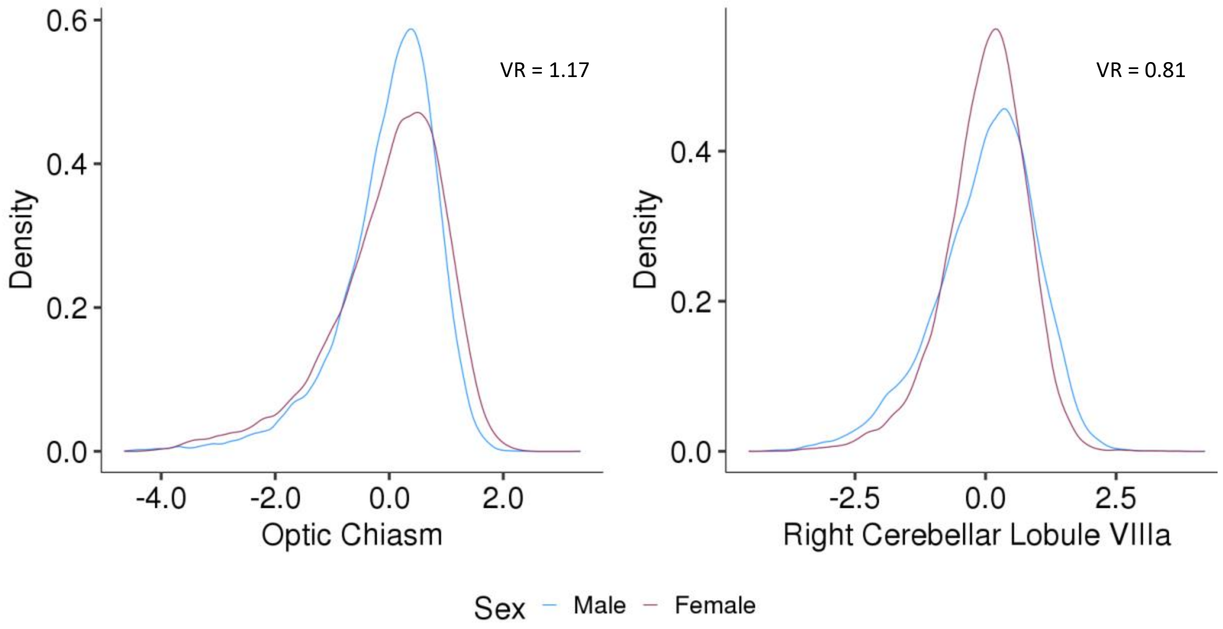


***Figure S6.*** *Largest Sex Differences in Variance Ratio across Brain Regions*. VR: Variance Ratio, Female SD / Male SD. Optic Chiasm was more variable in females and the Right Cerebellar Lobule VIIIa was more variable in males. No sex differences are seen as density is plotted against the residuals of the model with age, total brain volume, and sex as well as age$^2$, total brain volume, and sex interactions.

##### *6.1.1 Global Measures.*

The variance for log of the cerebellum GMV was significantly greater in males compared to females (VR = 0.90, F = 174.04, p = 1.18E-39) as well as the midbrain volume (VR = 0.97, F = 16.06, p = 6.16E-05).

##### *6.1.2 Cortical Volumes.*

Sex differences in variance were found for 46% of cortical volumes (68/148) ranging from 0.91 (right parahippocampal gyrus) to 1.14 (right subcallosal gyrus). Of the 68 regions, 5 were more variable in females than males, including the subcallosal gyri, right suborbital sulcus, right pericallosal sulcus, and right horizontal ramus of the anterior segment of the lateral sulcus volumes (Figure S7).

##### *6.1.3 Cortical Mean Thicknesses.*

Sex differences in variance were found for 42% of cortical volumes (62/148) ranging from 0.90 (left middle-anterior part of the cingulate gyrus and sulcus) to 1.15 (left anterior transverse collateral sulcus). More than half of the mean thicknesses exhibiting differences in variance between sexes were more variable in females than males (Figure S7).

##### *6.1.4 Cortical Surface Area.*

Sex differences in variance were found for 53% of cortical volumes (79/148) ranging from 0.90 (left opercular part of inferior frontal gyrus) to 1.14 (right subcallosal gyrus). Only 8 cortical surface areas were more variable in females than males: the subcallosal gyri, the lateral orbital sulci, the right suborbital sulcus, the left posterior transverse collateral sulcus, the lateral orbital sulcus, and the right inferior temporal sulcus (Figure S7).

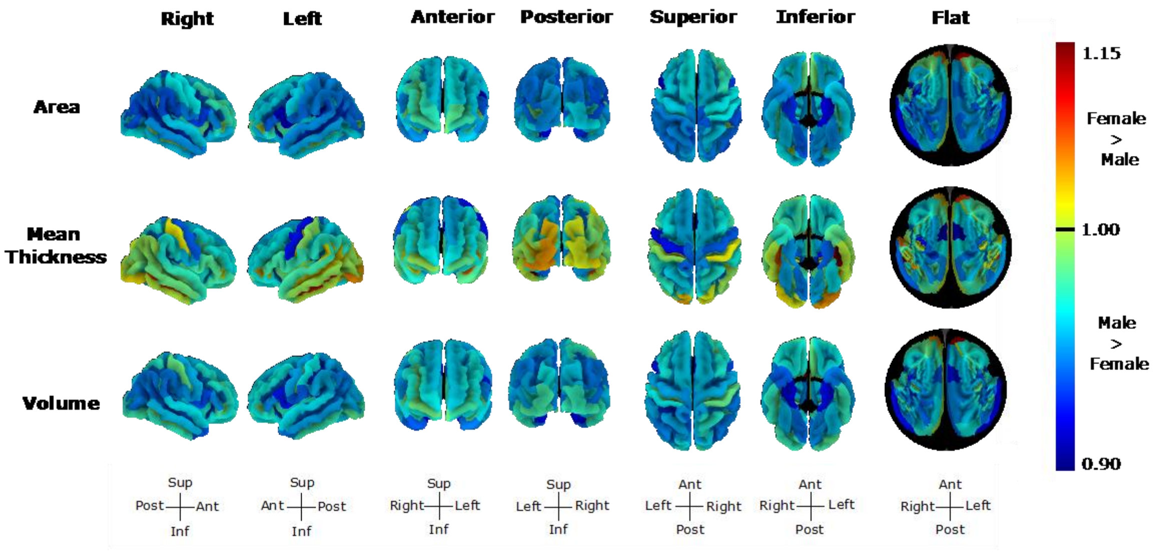


***Figure S7. Variance Sex Differences across Cortical Measures.*** Sex differences in variance (Variance Ratio = SD in females/SD in males) ranged from 0.90 (left surface area of the opercular part of the inferior frontal gyrus) to 1.15 (left mean thickness of the anterior transverse collateral sulcus). The flat representation corresponds to the flatten image of the superior view with the circle middle reflecting regions within the sagittal plane and circle edges reflecting inferior regions.

#### 6.1.5 ASEG Volumes.

A total of 14 out of 32 (44%) regions showed sex differences in variance ranging from 0.93 (posterior corpus callosum) to 1.17 (optic chiasm).

The left and right choroid plexus were more variable in females than males (VR= 1.11, VR = 1.07, respectively), while the 3rd and 4th ventricles were more variable in males than females (VR= 0.95, VR = 0.96, respectively). The left and right white matter volumes did not differ between sexes in terms of variance except for the optic chiasm volume, which was greater in females. Interestingly, the corpus callosum posterior volume variance was greater in males than females (VR= 0.93), while the corpus callosum central and mid-anterior volume variances were greater in females than males (VR= 1.04, VR= 1.02). Finally, the left and right caudate volumes which did not show sex differences in mean were more variable in males than females (VR= 0.95, VR= 0.96).

##### *6.1.6 Subcortical Freesurfer Volumes.*

Sex differences in variance were found in 64 out of 116 (55%) of whole subcortical and subsegmentations volumes and were in larger in males. The whole hippocampus (Left VR = 0.96, Right VR= 0.93) was the only whole subcortical volume that differed in terms of variance between sexes. The majority of hippocampal subsegmentations (28/40, 71%) were more variable in males than females, including regions such as the CA1, CA3, and CA4 bodies and heads, which differed in terms of mean sex differences. All hippocampal body volumes of the CA1, CA3, CA4, molecular layer, presubiculum, subiculum, and the granule cell layers of the dentate gyrus were more variable in males. About 60% (31/52) thalamic subsegmentations were more variable in males than females. The anteroventral regions and lateral geniculate nuclei which did not differ between sexes in terms of mean were more variable in males than females. Greater male variability was largely spread across thalamic regions (anterior, posterior, intralaminar, ventral, medial, and lateral regions).

##### *6.1.7 Cerebellar GMVs.*

About 89% of cerebellar regions exhibited sex differences in variance (25/28) ranging from 0.82 (right cerebellar lobule VIIIa) to 1.04 (left cerebellar lobule X). The left and right cerebellar lobule X were the only regions that were more variable in females than males (VR = 0.96, for both). The cerebellar vermis X, crus I vermis, and vermis VI did not vary differently between sexes.

### 6.2 Comparing TBV Adjustment Techniques – Allometric Versus Covariate or Proportion Adjustment

##### *6.2.1 Comparing Standardized Beta Coefficients when omitting and considering brain allometry.*

We examined result discrepancies between TCM adjustment techniques (allometric vs. linear covariate and allometric vs. proportion) across effects and interactions that were significant in our primary analyses. We did not look at differences in significance between TCM adjustment techniques for the TCM effect, since TCM was always a significant predictor of the brain region (except for the 5$^{th}$ ventricle). See Supplemental Table F1 for the consistency in the number of significant standardized beta coefficients across global measure adjustment techniques by segmentation.

We found conflicting results in 2.35% of statistical tests between the allometric and the linear covariate approach (102/4340). Discrepancies were reported in 3.44% of volumes (78/2268), 0.97% of surfaces (10/1036), and 1.35% of thicknesses (14/1036). There were more significant effects and interactions across cortical surface areas and mean thicknesses, subcortical subsegmentations, and cerebellar volumes in the models with the allometric adjustment compared to the models with the covariate adjustment. The opposite was true for ASEG and cortical volumes. Inconsistencies between allometric and linear covariate adjustment models were observed for effects and interactions near the threshold of significance (Supplemental Figures 3 https://osf.io/s4qc5/?view_only=7c6e633d701d443e96602a857cf75824).

Meanwhile, we found conflicting results in 14.24% of statistical tests between the allometric and the proportion approach (618/4340). Discrepancies were reported in 18.87 % of volumes (428/2268), 13.51% of surfaces (140/1036), and 4.83% of thicknesses (50/1036). The proportion method models yielded a greater number of significant effects and interactions across all cerebral measures compared to the models with the allometric adjustment.

The standardized beta varied across models, with a difference of up to 0.04 between the allometric and the covariate approach and up to 0.23 between the allometric and the proportion method (Supplemental Figures 3, https://osf.io/s4qc5/?view_only=7c6e633d701d443e96602a857cf75824). For instance, the standardized beta of the interaction of TBV by Sex reported for the right cerebellar VIIIa lobule was at -0.05 in the linear model and -0.09 in the allometric model and the standardized beta of the interaction of $Age^2$ by Sex reported for the right anterior pulvinar thalamic nucleus was at -0.23 in the linear model and 0.00 in the allometric model.

#### *6.2.2 Comparing the Sex Difference in Variance across TCM Adjustment Techniques.*

We found conflicting results in 38.06% of the reported sex differences in variance between the allometric and the linear covariate approach (236/620) and in 29.03% of the reported sex differences in variance between the allometric and the proportion approach (180/620). Discrepancies between the allometric and covariate adjustment models were reported in 46.91% of volumes (152/324), 46.62% of surfaces (69/148), and 10.14% of thicknesses (15/148), while discrepancies between the allometric and covariate adjustment models were reported in 37.04% of volumes (120/324), 30.41% of surfaces (45/148), and 10.14% of thicknesses (15/148). Moreover, 96 % of the sex differences in variance found in the allometric model were present in the linear model with the covariate adjustment for TBV and 37% for the linear model with the proportion adjustment for TBV. Finally, 57 % of the sex differences in variance found in the linear model with the covariate adjustment for TBV were present in the allometric model and 32% for the linear model with the proportion approach. Sex differences in variance were more common when the variance ratios were calculated from the proportion and linear covariate models than the allometric models across regional and global measures (see Supplemental Tables F2 and F3, respectively and Supplemental Figures 4 https://osf.io/s4qc5/?view_only=7c6e633d701d443e96602a857cf75824).

We also calculated the correlations between the standardized beta ($\beta < 0$: Female < Male) and the VR of the sex difference (VR < 1: Female < Male) to examine whether regions with the largest sex differences were also the most variable (Supplemental Figures 2 https://osf.io/s4qc5/?view_only=7c6e633d701d443e96602a857cf75824). In models with the linear covariate or proportion TCM adjustment, larger regions in females were more variable in females in subcortical ($r_{covariate} = 0.39$, $r_{proportion} = 0.48$), and ventricle, WM, and corpus callosum volumes ($r_{covariate} = 0.64$, $r_{proportion} = 0.71$). In contrast, larger regions in females were more variable in males ($\mu\ r = -0.33$) in the allometric adjustment for TCM and significant across cerebral regions except for the cerebellar and ventricle, WM, and corpus callosum volumes (Supplemental Table F4).

##### *6.2.3 Correlations between a region's deviance from isometry (i.e., |1 – scaling coefficient|) and the difference in standardized beta between the linear (covariate or proportion) and allometric models*

We examined the correlations between a region's deviance from isometry (i.e., |1 – scaling coefficient|) and the difference in standardized beta between the linear (covariate or proportion) and allometric models (e.g., |Covariate Standardized Beta – Allometric Standardized Beta|) to investigate whether the difference in standardized beta between the linear and allometric models occurred in regions with a greater degree of allometry.

Across all regions, the correlation between deviance from isometry and the difference in the standardized beta of all effects was 0.16 (p = 1.30e-13) for the allometric and the covariate models and was 0.28 (p = 2.22e-16) for the allometric and the proportion models.

When looking at the correlations by effect or interaction across regions (Supplemental Figures 5 https://osf.io/s4qc5/?view_only=7c6e633d701d443e96602a857cf75824), we found strong significant correlations between the deviance from isometry and the standardized beta difference of the allometric and proportion approach ranging from 0.41 (TBV x Sex) to 0.67 (TBV x Age) and smaller significant correlations for the standardized beta difference of the allometric and linear covariate approach ranging from 0.15 (Sex) to 0.28 ($Age^2$; Supplemental Table F5A). The strength and direction of the association varied across segmentations (Supplemental Table F5B).

#### 6.3 Replication of Sex Differences in Studies considering Brain Allometry

We attempted to replicate previous sex differences reported by studies taking into account brain allometry and found mixed results (see Supplemental Tables G1- 4).

In short, we replicated the sex differences in the frontal lobes described by Peyre and colleagues (2020) in children who did not adjust for age or $age^2$, as well as the greater pallidum volume in males reported by Reardon and colleagues' (2016). However, instead of being greater in males (Mankiw et al., 2017), we found that the cerebellum, flocculus, cerebellar lobule VIIb, VIIb, and VIIA, and Crus II volumes were greater in females and that the flocculus volume did not vary between sexes. Finally, Sanchis-Segura and colleagues (2019) adjusted for brain allometry using the power proportion method which does not take into account the intercept of the linear equation (log10(y) = Intercept + slope * log10(x)). We replicated only 10 out of the 25 mean sex differences identified by Sanchis-Segura and colleagues (2019) after an FDR correction for multiple comparison. It is important to note that the effect sizes in our sample and confidence intervals of our study were much smaller. The same significant results were reported when using the logarithmic equation to adjust for brain allometry, although effect sizes were greater and p values were smaller.

### 6.4 Replication of Ritchie and Colleagues (2018) Study and Comparing Results with the Covariate and with the Allometric TBV Adjustment.

We attempted to replicate the mean and variance sex differences reported by Ritchie and colleagues (2018) when adjusting brain volumes, thicknesses, and surface areas, for age, ethnicity, and total brain volume (for volumes) or total surface area (for areas) or total mean cortical thickness (for mean thicknesses). After providing a summary of the replication analyses and TCM adjustment comparison, we provided a detailed comparison by segmentation where we simultaneously discussed our replication results and compared our results from our replication analyses to those obtained when considering brain allometry. Replication and TCM comparison results are provided in Supplemental Tables G5-G9.

#### *6.4.1 Summary.*

The sex differences in mean and variance reported by Ritchie and colleagues (2018) in subcortical volumes from the FIRST segmentations and cortical volumes, mean thicknesses, and surface areas from the Desikan-Killiany Cortical Atlas were generally replicated, with some exceptions. More mean differences in volumes and surface areas and sex differences in variance for mean cortical thickness were reported in our replication compared to Ritchie and colleagues' (2018) study (Table 1).

Mean sex differences were consistent between the allometric and linear covariate TCM adjustments, except for the left pars opercularis and left transverse temporal volumes, whose mean sex differences were only significant when considering brain allometry. When taking into account brain allometry, the sex difference in variance was no longer significant for the left pallidum and the number of significant cortical sex differences in variance decreased in volumes and surface areas but increased in mean thicknesses (Table S1).

Finally, we replicated the absence of significant correlations reported by Ritchie and colleagues (2018) between the sex difference's Cohen's d and the VR in cortical and subcortical volumes, cortical mean thicknesses, and cortical surface areas. We found that when taking into account brain allometry, the correlation coefficients between the sex difference's Cohen's d and the VR in cortical and subcortical volumes decreased and became significant for cortical mean thickness (Supplemental Table G10).

Table S1. A Comparison of the Number of Significant Mean Sex Difference reported by Ritchie and colleagues (2018) to those reported in our Linear Covariate and Allometric Replications.

| | | **Ritchie and colleagues (2018)** | **Result Overlap** | **Present Study** | |
|---|---|---|---|---|---|
| | ***Total Brain Volume Adjustment*** | ***Linear Covariate*** | | ***Linear Covariate*** | ***Allometric*** |
| ***Significant Mean Sex Differences*** | Cortical Volumes | 23 | 19 | 54 | 56 |
| | Cortical Surface Areas | 25 | 24 | 41 | 41 |
| | Cortical Mean Thicknesses | 50 | 42 | 55 | 55 |
| | Subcortical Volumes | 6 | 6 | 11 | 11 |
| | Total N | 104 | 91 | 161 | 163 |
| | Total % | 52 | 45.5 | 80.5 | 81.5 |
| ***Significant Variance Sex Differences*** | Cortical Volumes | 58 | 58 | 62 | 55 |
| | Cortical Surface Areas | 59 | 59 | 62 | 56 |
| | Cortical Mean Thicknesses | 2 | 2 | 32 | 44 |
| | Subcortical Volumes | 14 | 14 | 14 | 12 |
| | Total N | 133 | 133 | 170 | 167 |
| | Total % | 66.5 | 66.5 | 85 | 83.5 |

*N.B.* Significance based on FDR correction. Result Overlap: Significant results reported both by Ritchie and colleagues (2018) and by our replication of their study with the linear covariate adjustment for Total Brain Volume. Total Cortical Measures from the Desikan-Killiany Atlas, N = 186 and from the Subcortical FIRST segmentations, N= 14.

***6.4.2 Subcortical Volumes: Mean and Variance Sex Differences.***

*Mean Sex Differences.* We replicated the majority of sex differences reported by Ritchie and colleagues (2018). However, in contrast with Ritchie and colleagues (2018) who did not find sex differences in the hippocampus, accumbens, and left thalamus volumes, we found that these regions were significantly greater in females compared to males in our replication when adjusting for TBV with the covariate and with the allometric approach.

*Variance Sex Differences.* We replicated Ritchie and colleagues' (2018) finding that males show greater variability across subcortical volumes when adjusting for TBV with the covariate approach. However, when adjusting for TBV with the allometric approach sex difference in variability for the left and right pallidum disappeared.

***6.4.3 Cortical Volumes: Mean and Variance Sex Differences.***

*Mean Sex Differences.* We replicated 19 of the 23 sex differences reported by Ritchie and colleagues. The right paracentral volume and the right superior temporal volume did not differ between sexes when adjusting for TBV with the covariate or the allometric approach, while sex differences in the left pars opercularis and left transverse temporal volume became significant after adjusting for brain allometry. We reported 54/62 significant sex when adjusting for TBV with the covariate approach and 56/62 when adjusting for brain allometry. Covariate and allometric approaches were consistent except for the left transverse temporal volume and left pars opercularis.

*Variance Sex Differences.* In addition to replicating the reported sex difference in volumetric variance by Ritchie and colleagues (2018), we additionally found greater male variability in the left pars orbitalis, left rostral anterior cingulate, and right pericalcarine volumes. However, after adjusting for brain allometry, the left caudal anterior cingulate, left isthmus cingulate, left and right medial orbitofrontal, left posterior cingulate, left insula, and right lateral orbitofrontal volumes no longer exhibited sex differences in variance.

***6.4.4 Cortical Surface Areas: Mean and Variance Sex Differences.***

*Mean Sex Differences.* We replicated 24 out of the 25 sex differences in mean reported by Ritchie and colleagues' (2018) when linearly adjusting for TSA, although we found that 41/62 regions differed between sexes. Ritchie and colleagues (2018) reported that females had a greater right supramarginal area, while the opposite was true in our analyses. Allometric and covariate approaches were consistent.

*Variance Sex Differences.* In comparison with Ritchie and colleagues (2018), we replicated all sex differences in variance and additionally reported significant effects in the right and left pericalcarine area and left pars orbitalis area. However, following adjustment for brain allometry, left caudal anterior cingulate, left medial orbitofrontal, left pericalcarine, right lateral orbitofrontal, right pericalcarine, and the right rostral anterior cingulate area no longer exhibited sex differences in variance.

***6.4.5 Cortical Mean Thicknesses: Mean and Variance Sex Differences.***

*Mean Sex Differences.* We replicated 42 out of the 50 significant sex differences reported by Ritchie and colleagues (2018). We did not replicate the sex differences in the left entorhinal, left pars orbitalis, left precentral, left precuneus, and right fusiform mean thicknesses when linearly adjusting for total mean cortical thickness or when taking into account brain allometry. Moreover, Ritchie and colleagues (2018) reported that the left lateral occipital, right inferior parietal, and right paracentral mean thickness were greater in males, while we found them to be greater in females. Finally, Ritchie and colleagues (2018) reported that the right inferior temporal mean thickness was greater in females, but the opposite was true in our analyses.

We found that 55/62 regions were significantly different between males and females. The left caudal middle frontal, left inferior temporal, left paracentral, left pars triangularis, right caudal middle frontal, right entorhinal, right lateral orbitofrontal, right pars triangularis, right rostral anterior cingulate, and right transverse temporal mean thicknesses did not significantly differ between sexes in Ritchie and Colleagues (2018) but did in the present study when considering the linear and allometric relationship of mean thicknesses with total mean cortical thickness.

*Variance Sex Differences.* In terms of sex differences in mean thickness variability, Ritchie and colleagues (2018) reported greater male variability in the left middle temporal mean thickness and greater female variability in the right inferior parietal mean thickness, while we found greater female variability in the left middle temporal mean thickness and no sex difference in variance in the right inferior parietal mean thickness. We found that 36/62 cortical mean thicknesses differed between sexes in terms of variability when linearly adjusting for total mean cortical thickness and 44/62 when considering brain allometry. The left lingual, left pericalcarine, and left post-central regions were no longer significant when considering brain allometry, while an additional 10 other regions significantly differed in variance between sexes.

### 6.5 Height

To investigate whether height explains part of the observed main effects and interactions, we ran the models with height instead of TBV and found that height was a significant positive predictor of all global measures except for total mean cortical thickness and the interaction of sex x log10(height) was significant for Cerebellar GMV ($\beta = -0.07$) at the strictest threshold (0.05/ (9*11)); Supplemental Table B8). When adding TBV as an interactive term, both log10(TBV) x sex ($\beta = -0.09$) and log10(height) x sex ($\beta = -0.07$) were significant at $p < (0.05/(9*23))$. Adding height in the model increased the β of the log10(TBV) x sex from -0.06 to -0.09. Thus, our findings suggest that the variance attributed to the log10(TBV) x sex interaction differs from the variance attributed to log10(height) x sex.

### 7- R packages

The following R packages were used in the study:

- data.table (Dowle et al., 2020)
- ggplot2 (Wickham, Chang, et al., 2020)
- tidyr (Wickham & RStudio, 2020)
- broom (Robinson et al., 2020)
- sjPLot (Lüdecke et al., 2020)
- car (Fox et al., 2020)
- ggpubr (Kassambara, 2020)
- plyr (Wickham, 2020)
- dplyr (Wickham, François, et al., 2020)
- MatchIt (Ho et al., 2011)
- Tables (Murdoch, 2020)
- Scales (Wickham, Seidel, et al., 2020)
- Corrplot (Wei et al., 2017)
- Ggrepel (Slowikowski et al., 2020)
